# Genome assemblies of Indian *desi* cattle reveals hotspots of rearrangements and immune-related genetic diversity

**DOI:** 10.1101/2025.01.14.632934

**Authors:** Sarwar Azam, Abhisek Sahu, Mohammad Kadivella, Aamir Waseem Khan, Mahesh Neupane, Curtis P Van Tassell, Benjamin D Rosen, Ravi Kumar Gandham, Subha Narayan Rath, Subeer S Majumdar

## Abstract

India, home to the world’s largest cattle population, hosts native dairy breeds essential to its agricultural economy because of their adaptability and resilience. This study characterizes the genomes of five prominent breeds Gir, Kankrej, Red Sindhi, Sahiwal and Tharparkar, highlighting their unique genomic characteristics. The *de novo* assemblies ranged from 2.70-2.78 Gb in size, with 90% of the genomes assembled in just 56 to 1,663 scaffolds. The use of reference-guided scaffolding further enhanced these genomes, resulting in 93.3-96.7% pseudomolecule coverage with strong BUSCO scores (94.1-95.5%). Comparative analyses revealed 87–95% synteny with the Brahman genome and identified 19.84–153.16 Mb of structural rearrangements per genome, including inversions, translocations, and duplications. Synteny diversity analysis uncovered 10,643 perfectly collinear regions spanning 87.3 Mb and 6,622 hotspots of rearrangement (HOT regions) covering 55.18 Mb. These HOT regions, characterized by high synteny diversity, were significantly enriched with immune-related genes. Moreover, immune-related gene clusters, including MHC, NKC, and LRC, were identified within HOT regions in the *desi* reference genome. Our findings provide valuable insights into the genetic diversity of *desi* cattle breeds. The high-quality genome assemblies generated in this study will serve as valuable resources for future research in genetic improvement, disease resistance, and environmental adaptation.

## Introduction

India is home to the largest cattle population in the world, with over 192 million head, including 50 recognized native zebu breeds (https://www.data.gov.in/catalog/20th-livestock-census) (1). Both indigenous *Bos indicus* and non-native *Bos taurus* breeds of cattle, as well as their crossbreeds, can be found throughout the country. Through traditional animal husbandry practices, these native, or *desi,* cattle have evolved over centuries to adapt to their harsh environments, developing resistance to tropical diseases and parasites, and the ability to subsist on poor quality roughages and grasses (1). The diverse range of agro-ecological zones in India has contributed to the development of a wide variety of cattle populations. The State of the World’s Animal Genetic Resources recognizes 60 local, eight regional transboundary, and seven international transboundary cow breeds from India (2). Among the highest milk producing Indian breeds are Gir, Kankrej, Red Sindhi, Sahiwal, and Tharparkar (3, 4). These breeds are native to India’s north-western region and correspond to three nearby agroclimatic zones. The Tharparkar breed is found in the western dry region (5), while Gir and Kankrej are located in the Gujarat hills and plains (6–8). Sahiwal, originating in the Sahiwal district of Pakistan near the Indo-Pak border, is primarily found in the trans-Gangetic plains (9) and Red Sindhi originated in Sindh, Pakistan, and is associated with the western dry region of India (10). Recent Indian livestock surveys have shown an increase in crossbreeding of these breeds (https://www.data.gov.in/dataset-group-name/19th%20Livestock%20Census). The government and non-profit organizations are actively promoting these breeds to maintain and enhance India’s milk output (9).

Breeds within a species are very similar, yet they exhibit differences in phenotypic characteristics such as appearance, adaptation, and behavior. Since phenotype is an expression of genotype, these variations can be captured at the genomic level through comparative genomics, particularly in identifying syntenic and non-syntenic or rearranged regions within the genome (11–14). Chromosomal rearrangements are typically studied in terms of structural variations (SVs) and sequence variations (15–17). Structural variations represent large genomic differences, such as large insertions, deletions, inversions, duplication, and translocations, while sequence variations represent smaller genomic differences, including single nucleotide polymorphisms (SNPs) and short insertions or deletions, copy loss, and copy gain within larger blocks of SVs or syntenic regions. SVs have recently been studied in various animal and plant species where multiple chromosomal-level genome assemblies are available (17–21). Several SVs have been identified as associated with key phenotypic traits in diverse plants and animals, including heat tolerance in pearl millet (22), nematode resistance in soybean (23), reproductive morphology in cucumber (24), coat color patterns in European domestic pigs (25), tick resistance in indicine cattle (26), color sidedness in cattle (27), and white coat color in sheep (28).

However, a basic requirement for synteny and re-arrangement studies is the availability of chromosomal-level assemblies of multiple individuals of the species (17, 29). Generating these assemblies is a tedious process, but with the availability of long-read sequencing technologies, it has become feasible to generate multiple chromosome-level assemblies for a species. The cattle genome was first published in 2009 (30), and since then, a few chromosomal-level assemblies of cattle have been developed. Most of these assemblies belong to the *Bos taurus* species, with the exception of a Brahman genome assembly (31), which represents the Brahman breed of *Bos indicus*. At the initiation of this study, no other *Bos indicus* chromosomal-level genome assemblies were publicly available. Therefore, we undertook the task of characterizing the genomes of the prominent dairy breeds of India using linked-read technology (32, 33). Linked reads, a technology that combines the advantages of short reads with the features of long reads, have been shown to provide chromosomal-level assemblies in humans and other species (34–38).

Despite the importance of synteny or collinearity (17), there is still little known about the actual degree of collinearity within populations, as most current genome studies are not based on chromosome-level assemblies. In this study, we aim to characterize the genomes of the five prominent Indian dairy cattle breeds: Gir, Kankrej, Red Sindhi, Sahiwal and Tharparkar using linked-read technology. We also analyzed the syntenic and rearranged regions among these breeds, quantified synteny diversity in *desi* cattle, and catalog collinear regions as well as hotspots of genomic rearrangement. Furthermore, we investigated the gene content and functional implications of these regions.

## Materials and methods

### Genome sequencing of cattle breeds

Blood samples were collected from representative individuals of the Gir, Kankrej, Red Sindhi, Sahiwal, and Tharparkar breeds, in compliance with the guidelines of the Committee for the Purpose of Control and Supervision on Experiments on Animals (CPCSEA), India, with the Institutional Animal Ethics Committee (IAEC) approval. High molecular weight (HMW) DNA was extracted by AgriGenome Pvt. Ltd. (Genome Sequencing Company, Bangalore, India), and 10X Genomics Chromium technology (39) libraries were prepared. Sequencing was performed on an Illumina HiSeq X system.

### Generation of *de novo* assemblies

Genomes of each breed of cattle were assembled with Supernova Assembler v2.1.1 (39) using 10X linked-reads as input. The Supernova assembler was provided with paired-end raw reads derived from the sequencing using option “run” with --maxreads. The output was generated using Supernova “mkoutput” and saved in “pseudohap” format as raw Supernova genome assemblies. Sequences composed entirely of N were removed. ARCS v1.2.4 (40)was used to further improve the assembly in --arks mode, which uses exact K-mer mapping and associates the barcode information of linked reads to order and orient the sequences. The ARCS “--arks” mode was run using default parameters on barcoded interleaved FASTQ data from LongRanger v2.2.2 (33) with raw Supernova assemblies to create the longer scaffolded assemblies. ARCS output was further processed using the GapCloser program from SOAPdenovo2 (41). Each output sequence was considered a *de novo* genome assembly, and the whole process was repeated five times to assemble each of the breed-specific genomes.

### Reference-guided assembly and pseudomolecule construction

*De novo* genome assemblies of each breed of cattle were further scaffolded using a template of Brahman genome assembly. The Brahman genome assembly (UOA_Brahman_1) (31) was obtained from NCBI and used as a reference genome. RagTag v2.1.0 (42), which is a collection of software for scaffolding and improving genome assemblies, was used on default parameters with the option “--scaffold” to generate reference-guided assemblies.

### Assembly completeness assessment

The completeness of assemblies in terms of gene space was assessed by the Benchmarking Universal Single-Copy Orthologs (BUSCO) tool (43). The BUSCO analysis was performed on *de novo* and reference-guided assemblies using the “mammalia_odb10” database with parameter -m “genome” using BUSCO v5.0.3. Outputs were produced in text format, which summarizes the BUSCO gene annotation.

### Assembly annotation

Each newly assembled reference-guided genome underwent structural annotation, encompassing gene annotation and repeat annotation. Gene annotation was carried out using Liftoff v1.6.3 [48]. Liftoff is a specialized tool designed to efficiently transfer gene annotations from a well-annotated reference genome to a newly assembled genome of a closely related species. The Brahman genome served as the reference, and its gene annotations were lifted over to the target genomes. The resulting annotations were saved in GFF3 format. A Perl script was used to summarize the annotations, and shared protein-coding genes across the genomes were visualized using a Venn diagram [49].

To identify repetitive elements and transposable elements (TEs), the genomes of each breed were analyzed using RepeatMasker v4.1.0. RepeatMasker was executed with the following parameters: --species cow, -xsmall, and -nolow.

### Genomic synteny and rearrangement analysis

Pseudomolecules generated using reference-guided assemblies were compared with the Brahman reference genome for structural variation identification and synteny analysis. This comparison required a multiple-step analysis in which both the reference and the query genomes were masked for repeats using RepeatMasker v4.1.0. Masked query chromosomes (pseudomolecules) were aligned against the masked reference chromosomes using the nucmer program of the mummer v4.0.0 package (44) which generates a delta file. Alignments were further processed using the “delta-filter” and “show-coords” programs of mummer. Synteny and Rearrangement Identifier (SyRI) was used for identifying inter and intra chromosomal structural rearrangements. SyRI v1.6.3 (45) was run with repeat masked reference and query chromosomes, an aligned coordinate file, and a delta file generated from mummer. The SyRI software provided results in TSV and VCF formats. Structural rearrangements between the genomes were visualized using plotsr v0.5.4 (46). Furthermore, we investigated the variation in SNP and short indel density within syntenic and rearranged regions.

### Quantification of genome collinearity and synteny diversity analysis

To quantify the degree of collinearity between assembled genomes, we employed synteny diversity (π_syn_), a parameter previously defined by Jiao et al. (_17_). Synteny diversity represents the average proportion of non-syntenic sites across pairwise genome comparisons within a population. To calculate π_syn_, all assembled genomes (n=5) and the Brahman reference were subjected to pairwise alignments using SyRI. Perl scripts were used to extract syntenic blocks from the SyRI output, which served as the basis for π_syn_ calculation. Synteny diversity (π_syn_) was calculated across the entire genome using a 5 kb sliding window with a step size of 1 kb. The π_syn_ values range from 0 to 1, with higher values indicating a greater proportion of non-collinear regions between the genomes. To visualize collinear and non-collinear regions in *desi* cattle, the calculated π_syn_ values were plotted using R. We extracted genes from collinear regions and compared them to a set of 9,226 conserved orthologous genes from the BUSCO “mammalia_odb10” database (43). This comparison allowed us to assess the evolutionary conservation of these genes across mammals.

### Hotspots of rearrangements analysis

Regions exhibiting a synteny diversity > 0.5 were designated as hotspots of rearrangements (HOT regions) (17), characterized by multiple accessions containing diverse haplotypes. To identify these regions, we examined π_syn_ in 5-kb sliding windows with a 1-kb step size across the entire genome. Adjacent HOT regions separated by less than 2 kb were merged.

To gain deeper insights into the characteristics of HOT regions, we conducted a comparative analysis of gene and TE content between these HOT regions and syntenic regions. This analysis aimed to characterize associations between genomic features and rearrangement hotspots.

### Gene Ontology and Quantitative Trait Locus analysis of genes in HOT regions

To explore the functional properties of genes located within HOT regions, we performed GO enrichment analysis using the ‘clusterProfiler’ R package (47) from Bioconductor. A significance threshold of *P-*value ≤ 0.05 (FDR by Benjamini-Hochberg) was applied to identify significantly enriched GO terms within the *Bos indicus* (Brahman) reference annotation (https://github.com/ASBioinfo/Genome_wide_annotation_Bos_indicus). The 10 most significant terms in each GO category (biological process, molecular function, and cellular component) were then visualized using a dotplot and barplot. We also investigated potential links between genes in HOT regions and major cattle quantitative trait loci (QTLs). We downloaded the annotation of the ARS_UCD 1.2 genome assembly (48) and performed a BLASTP search against the annotated proteins using the genes present in HOT regions. Identified orthologous genes were then cross-referenced with a publicly available cattle QTL database (https://www.animalgenome.org/cgi-bin/QTLdb/BT/index). This analysis aimed to identify enriched QTLs and their associated traits.

### Characterization of immune gene clusters and synteny diversity

To investigate the impact of synteny diversity within immune gene clusters (IGCs) of *desi* cattle, we identified homologous regions in the Brahman genome corresponding to previously reported *Bos taurus* IGC locations (49). The *Bos taurus* genome assembly UCD-ARS1.2 has been well-characterized for all three major IGC regions: the major histocompatibility complex (MHC) on Chr 23 (28.3–28.7 Mb), the natural killer complex (NKC) on Chr 5 (99.5–99.8 Mb), and the leukocyte receptor complex (LRC) on Chr 18 (63.1–63.4 Mb). We conducted BLASTN (50) searches using known *Bos taurus* IGC coordinates to identify IGC regions in the Brahman genome. The boundaries of each IGC region were further refined and plotted using Gepard (51). Subsequently, we scanned these IGC regions for synteny diversity (π_syn_) values calculated earlier. To analyze orthologous genes within IGC regions across the breeds, we clustered annotated genes from each breed using OrthoFinder (52). By integrating orthologous gene clusters, alignment data, and IGC region coordinates on the Brahman genome, we generated a comprehensive gene block diagram visualizing syntenic relationships and gene order for each IGC cluster using Perl scripts.

## Results

### *De novo* genome assembly of five indigenous cattle breeds

To characterize and assemble the genomes of indigenous cattle breeds, we sequenced five breeds (Gir, Kankrej, Red Sindhi, Sahiwal and Tharparkar) using 10X Chromium technology. Each genome was sequenced to approximately 100x coverage with 150bp paired-end reads. *De novo* genome assemblies were generated in two steps: First, linked reads were assembled using the Supernova assembler **(Supplementary Table S1)**. Second, these assemblies were merged and polished using ARCS and Gapcloser (**Supplementary Figure S1**). The resulting *de novo* assemblies ranged from 2.70 to 2.78 Gb, closely resembling the Brahman genome assembly (2.7 Gb), which is publicly available as a reference genome for *Bos indicus* species. These assemblies exhibited high contiguity, with 90% of each genome being assembled in 56, 86, 817, 1,443, and 1,633 scaffolds for Sahiwal, Red Sindhi, Kankrej, Gir, and Tharparkar, respectively **(Table 1)**. However, the N50 scaffold lengths of Sahiwal and Red Sindhi assemblies were significantly longer (65.7 Mb and 45.8 Mb, respectively) than those of Gir, Kankrej, and Tharparkar. These results indicate Sahiwal and Red Sindhi are superior assemblies. Interestingly, many scaffolds in Sahiwal and Red Sindhi assemblies approached chromosomal length, suggesting they were near pseudomolecule-level assembly. The completeness of these assemblies was assessed using BUSCO analysis, which revealed that all *de novo* assemblies captured at least 93.3% of BUSCOs, indicating a high level of genomic completeness **(Figure 1A)**. Red Sindhi and Sahiwal had the highest BUSCO completeness (95.4%), while Tharparkar had the lowest (93.3%). These results were comparable to the reference Brahman assembly, which had a BUSCO annotation rate of 95.7%. Overall, the BUSCO analysis confirmed the high quality and completeness of these *de novo* assemblies.

**Figure 1:**
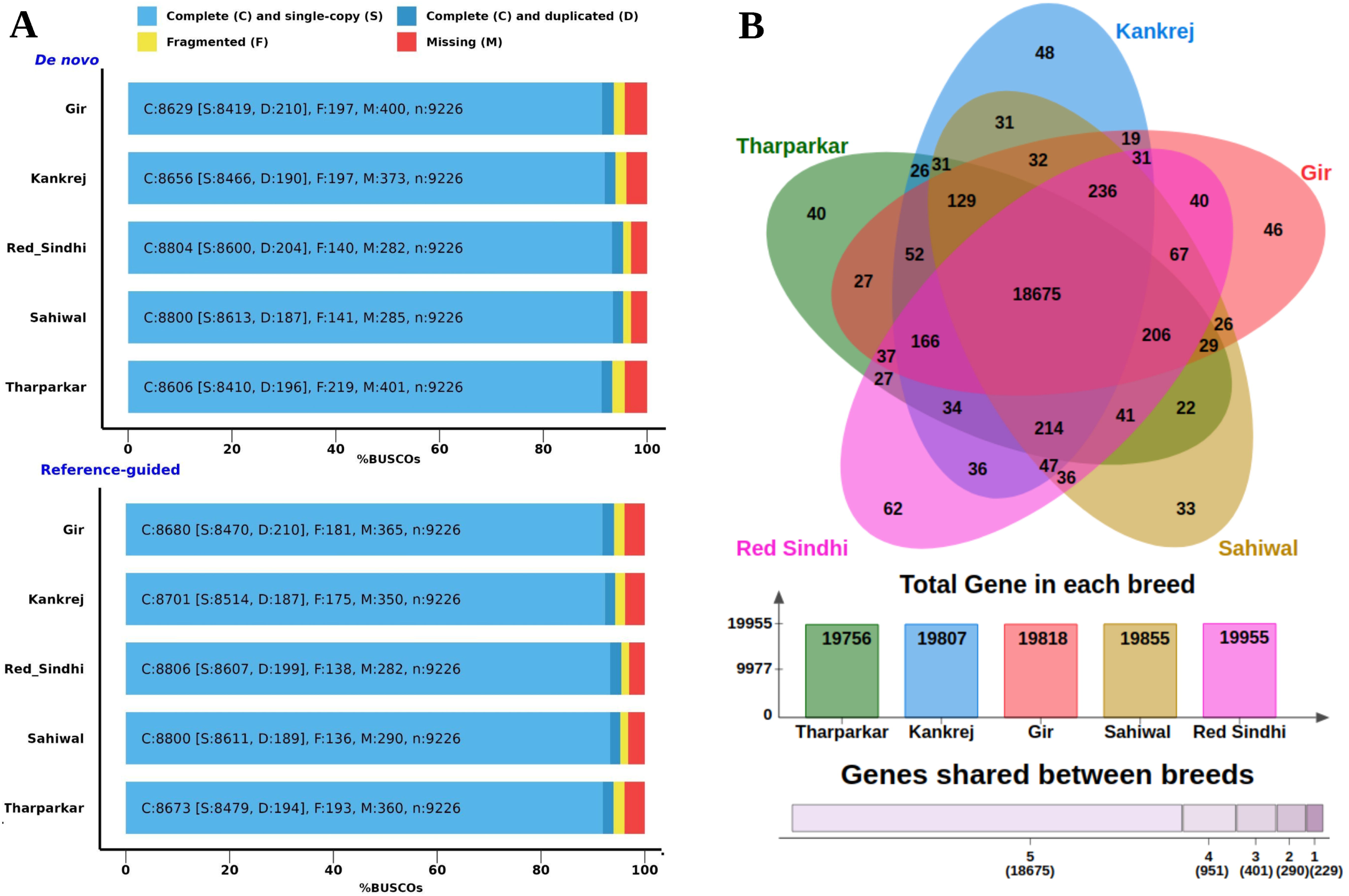
Liftoff annotation and completeness of the *desi* cattle genome. **(A) Busco analysis of genome assemblies.** The genome assembly and gene set were assessed for completeness using BUSCO. All *de novo* assemblies (upper panel) achieved a completeness of over 93%, while the reference-guided assembly (lower panel) reached 94%. **(B) Gene annotation in genome assemblies.** Venn diagram illustrates the proportion of genes lifted from the Brahman genome assembly using Liftoff for each *desi* breed.

**Table 1:** Statistics of *desi* cattle *de novo* genome assembly.

| **Genomic feature** | **Gir** | **Kankrej** | **Red Sindhi** | **Sahiwal** | **Tharparkar** |
| --- | --- | --- | --- | --- | --- |
| **No. of Scaffolds** | 35,073 | 28,895 | 20,404 | 21,876 | 38,144 |
| **Assembly size (bp)** | 2,773,546,526 | 2,732,556,869 | 2,701,116,505 | 2,713,754,657 | 2,780,671,120 |
| **Largest Scaffold (bp)** | 13,422,926 | 55,744,479 | 151,580,471 | 156,124,625 | 18,339,566 |
| **Smallest Scaffold (bp)** | 1,000 | 1,000 | 973 | 987 | 968 |
| **N50 (bp)** | 2,601,999 | 4,378,674 | 45,763,463 | 65,662,462 | 2,543,532 |
| **L50 (No.)** | 309 | 153 | 17 | 15 | 319 |
| **N90 (bp)** | 175,889 | 366,322 | 4,931,469 | 6,770,838 | 143,200 |
| **L90 (No.)** | 1,443 | 817 | 86 | 56 | 1,633 |

### Generation of reference-guided assemblies

Draft genome assemblies can be further improved to chromosome level using various strategies like scaffolding with long reads, Hi-C data, linkage map, optical map, etc. Another way of improving the draft genome to chromosome or pseudomolecules level assembly is to use a reference-guided approach, where the scaffolder uses a closely related genome as template. Draft *de novo* assemblies of each breed were further clustered, ordered, and oriented into pseudomolecules by a reference-guided approach using the Brahman genome assembly as the template. The results showed that 93.3% to 96.7% of the genome was assembled into pseudomolecules, with the highest percentage for Sahiwal (96.71%) and Red Sindhi (96.58%), and the lowest for Gir (93.38%) (**Figure 2; Supplementary Table S2**). The small portion of the genome assembly that was not merged into pseudomolecules represented a high number of contigs that were quite short and fragmented. In comparison to the Brahman reference genome, which had 1,250 total sequences and 97.88% of its bases assembled into pseudomolecules, the number of scaffolds in each assembly was higher, with Tharparkar, Gir, Kankrej, Red Sindhi, and Sahiwal having 27,311, 25,313, 20,861, 16,082, and 17,113, respectively. However, the N50, L50, and other assembly statistics were similar to the Brahman reference genome, indicating that high-quality reference-guided assemblies of each breed were achieved. The reference-guided approach also improved the BUSCO statistics with overall annotation rates ranging from 94.0% to 95.5% **(Figure 1A; Supplementary Table S3)**. These more contiguous and complete assemblies have been used for all downstream analyses including gene annotation, structural variation, and comparative genomics.

**Figure 2:**
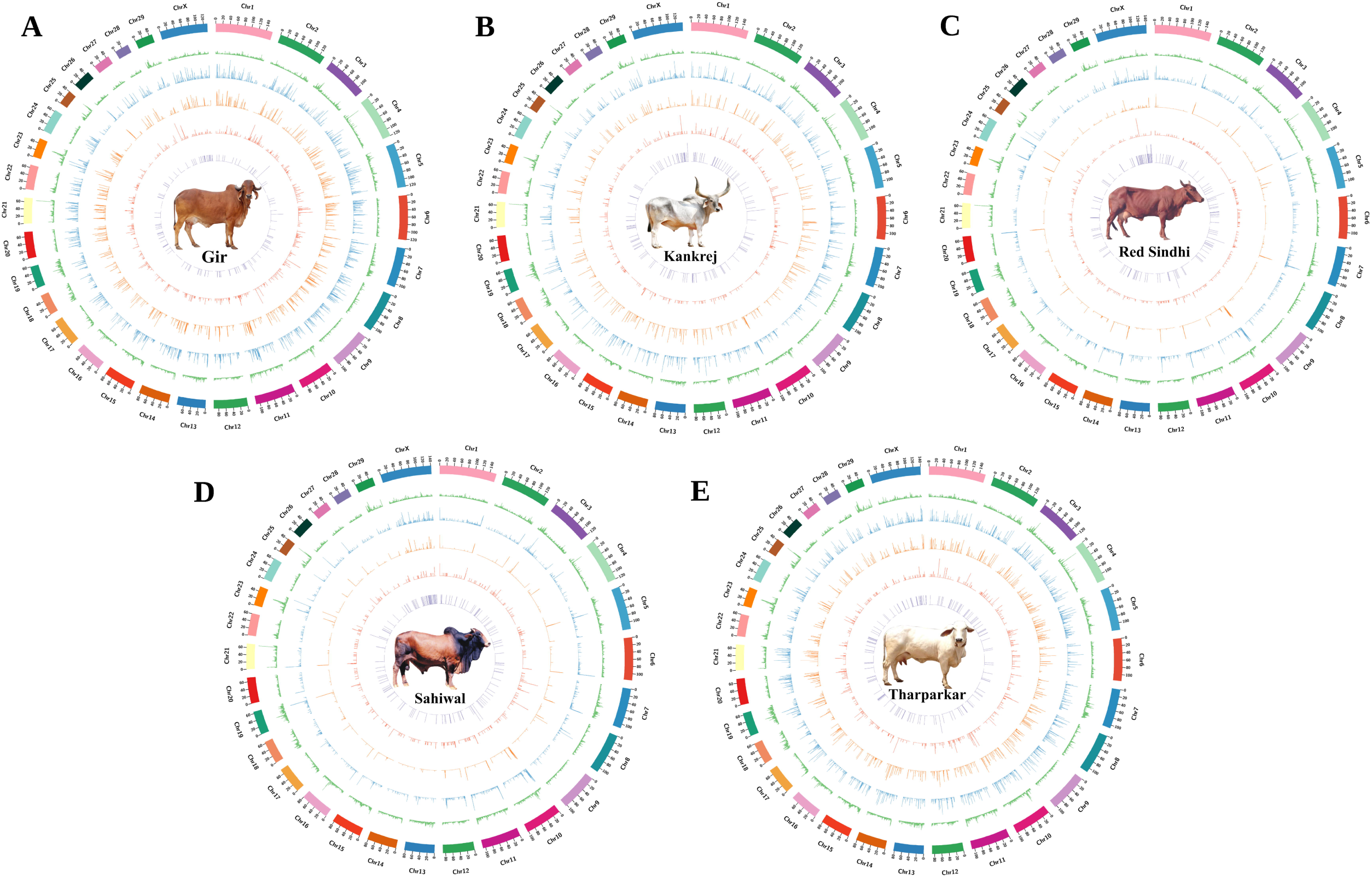
Genomic variations in desi cattle assemblies. The Circos plot provides a comprehensive visualization of the genomic architecture of five Indian cattle breeds: (A) Gir (B) Kankrej (C) Red Sindhi (D) Sahiwal (E) Tharparkar. The outermost track represents the chromosomes scaled to their respective sizes. Moving inward, the subsequent tracks include gene density (green), genome-specific sequence density (blue), translocation density (orange), duplication density (pink), and inversion density (violet). Density of various genomic features were calculated within non-overlapping 100 kb windows across the genome.

### Liftover genes in assemblies

Gene annotation for each of the five *desi* breeds was conducted by transferring annotations from the reference Brahman genome. The Brahman genome contains 28,038 annotated genes, of which the majority (20,846) were successfully transferred to the *desi* breed genomes. The number of transferred protein-coding genes varied slightly among the breeds, with Red Sindhi having the largest number and Tharparkar having the smallest. However, the overall differences were minimal, and the distribution of these genes revealed a high fraction of shared genes and breed-specific genes. In total, 18,675 genes were found to be common across all five breeds. Additionally, each breed exhibited unique gene content, with Sahiwal, Tharparkar, Gir, Kankrej, and Red Sindhi having 33, 40, 46, 48, and 62 unique genes, respectively (**Figure 1B**). Beyond protein-coding genes, various RNA types, including rRNA, tRNA, lncRNA, guide RNA, snoRNA, and snRNA, were also identified across the genomes (**Supplementary Table S4**).

### Repetitive element composition in genome assemblies

Each assembled genome was annotated for various types of repetitive elements **(Supplementary Table S5)**. The distribution of repeats across the assemblies was consistent and comparable to that of the Brahman reference genome. Analysis revealed that *desi* cattle breeds have an estimated ∼47% of their genomes composed of repetitive elements. Long interspersed nuclear elements (LINEs) were the most abundant class of repeats (∼27%), followed by short interspersed nuclear elements (SINEs), long terminal repeats (LTRs), and DNA elements. Only a small fraction (0.02%) of interspersed repeats remained unclassified.

### Syntenic and rearranged regions in *desi* cattle

To assess genomic architecture, we compared each assembled genome to the Brahman reference. Syntenic, rearranged (inversions, duplications, translocations), and genome-specific regions were identified through whole-genome alignment (**Figure 3A**). A range of 87-95% of each genome exhibited synteny with the reference, with Red Sindhi displaying the highest and Gir the lowest synteny. Rearranged regions totalled 19.8-153.2 Mb per genome, comprising 2.6-5.3 Mb of inversions (127–147), 1.8-6.4 Mb of polymorphic duplications (970–1,572), and 14.9-144.0 Mb of translocations (1,717–10,111) (**Figure 3B; Supplementary Table S6**). These structural rearrangements (inversions, duplications, translocations) within each genome vary in size, ranging from hundreds of base pairs to megabases (**Figure 3C; Additional file 1**). Translocations were the most abundant structural variations, with the majority being interchromosomal **(Table 2)**. In contrast, inversions tended to be larger than other structural rearrangements, with many exceeding 100 kb **(Supplementary Table S7)**. The largest inversion, measuring 2.8 Mb on Chromosome X, was found in Red Sindhi (**Figure 4A**). Notably, a large inversion on chromosome 12 was found in all the *desi* genomes compared to the Brahman reference. This sequence contains lncRNA, SnRNA, and a protein-coding gene, LMO7.

**Figure 3:**
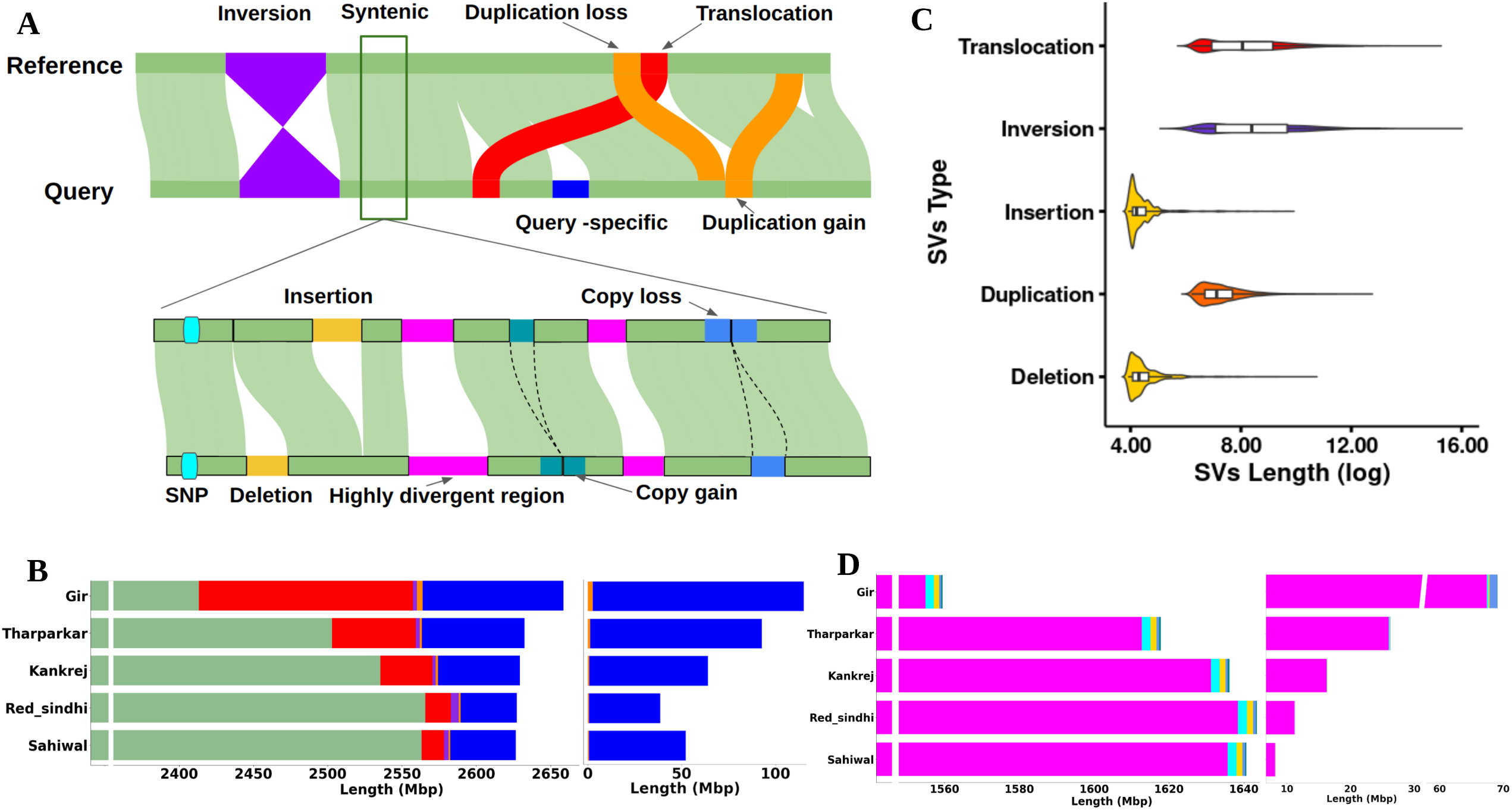
Syntenic and rearranged regions between genomes. **(A)** Schematic representation of structural differences (upper panel) and sequence variations (lower panel) identifiable between genome assemblies. Note that local sequence variation can occur in both syntenic and rearranged regions. **(B)** The bar plots on the left show the total span of syntenic and rearranged regions between the reference genome and each of the other accessions, while the right plot shows the sequence space specific to each accession (colors correspond to the schematic diagram of structural differences in (A)). **(C)** Length distributions of different types of major structural and sequence variations in the genome. **(D)** The bar plots display local sequence variation in syntenic (left) and rearranged (right) regions between the reference genome and each of the other accessions (colors correspond to the schematic diagram of sequence differences in (A)).

**Figure 4:**
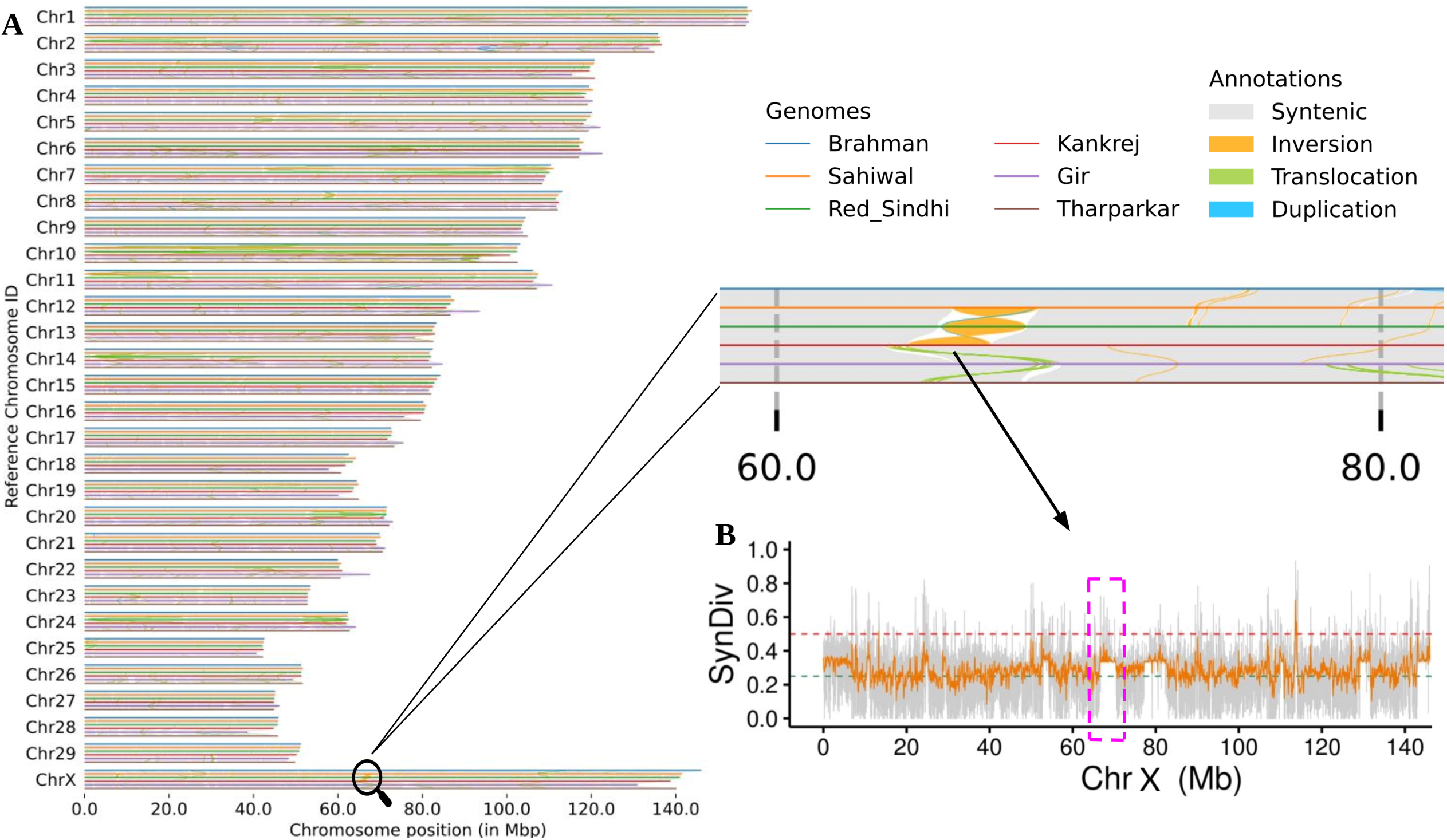
A detailed view of the largest inversion in the Red Sindhi on chromosome X. **(A)** The left panel shows a synteny and rearrangement plot across all chromosomes for six major cattle breeds including Brahman, with a zoomed view highlighting the 2.83 Mb inversion on Chromosome X. **(B)** Synteny diversity across chromosome X is depicted with two sliding window sizes: 100 kb windows with a 50 kb step (in orange), and 5 kb windows with a 1 kb step (in grey). The arrow points to the region (dashed pink line) where 2.83 Mb inversion observed in the Red Sindhi.

**Table 2:** Number and total length of rearranged regions in *desi* cattle genomes.

| Sample Name | INV | ITX | CTX | DUP-Loss | Ref. Specific | DUP-Gain | Acc. Specific |
| --- | --- | --- | --- | --- | --- | --- | --- |
| *Number* |  |  |  |  |  |  |  |
| **Gir** | 134 | 2,628 | 7,483 | 815 | 11,248 | 757 | 11,771 |
| **Kankrej** | 127 | 3,865 | 33 | 641 | 6,105 | 400 | 5,967 |
| **Tharparkar** | 146 | 5,405 | 275 | 623 | 7,960 | 545 | 8,096 |
| **Sahiwal** | 147 | 1,899 | 23 | 609 | 4,133 | 395 | 3,886 |
| **Red Sindhi** | 145 | 1,695 | 22 | 615 | 3,731 | 355 | 3,398 |
| *Length (bp)* |  |  |  |  |  |  |  |
| **Gir** | 2,688,837 | 25,663,618 | 118,344,504 | 3,824,712 | 94,802,956 | 2,643,892 | 112,459,228 |
| **Kankrej** | 2,298,347 | 34,894,573 | 115,407 | 1,301,825 | 55,178,141 | 859,024 | 63,290,939 |
| **Tharparkar** | 2,647,949 | 53,337,548 | 2,879,661 | 1,467,263 | 69,070,685 | 1,278,720 | 91,443,475 |
| **Sahiwal** | 3,125,028 | 14,838,983 | 76,628 | 1,133,576 | 44,067,767 | 675,218 | 51,645,093 |
| **Red Sindhi** | 5,317,975 | 17,012,560 | 99,134 | 1,360,229 | 1,360,229 | 577,108 | 38,188,257 |

**Note:** **INV:** inversion; **ITX:** intra-chromosome translocation; **CTX:** inter-chromosome translocation; **DUP-Loss:** duplication loss in accession; **Ref. Specific:** regions specific to the reference genome; **DUP-Gain:** duplication gain in accession; **Acc. Specific:** regions specific to the accession.

Further analysis of alignments revealed sequence variations like SNP, highly divergent region, copy gain, copy loss and InDels. Each genome contained over 2 million SNPs, with Red Sindhi exhibiting the highest number followed by Sahiwal (**Figure 3D; Supplementary Table S8**). Sequence copy number variation analysis identified 113-218 copy gain events per genome, with Tharparkar showing the highest number, primarily located in syntenic regions. Conversely, copy loss events were more abundant and larger in size across all genomes except Gir. Sequence variation analysis revealed a higher density of SNPs (**Figure 5A)** and INDELs (**Figure 5B**) within rearranged regions compared to syntenic regions.

**Figure 5:**
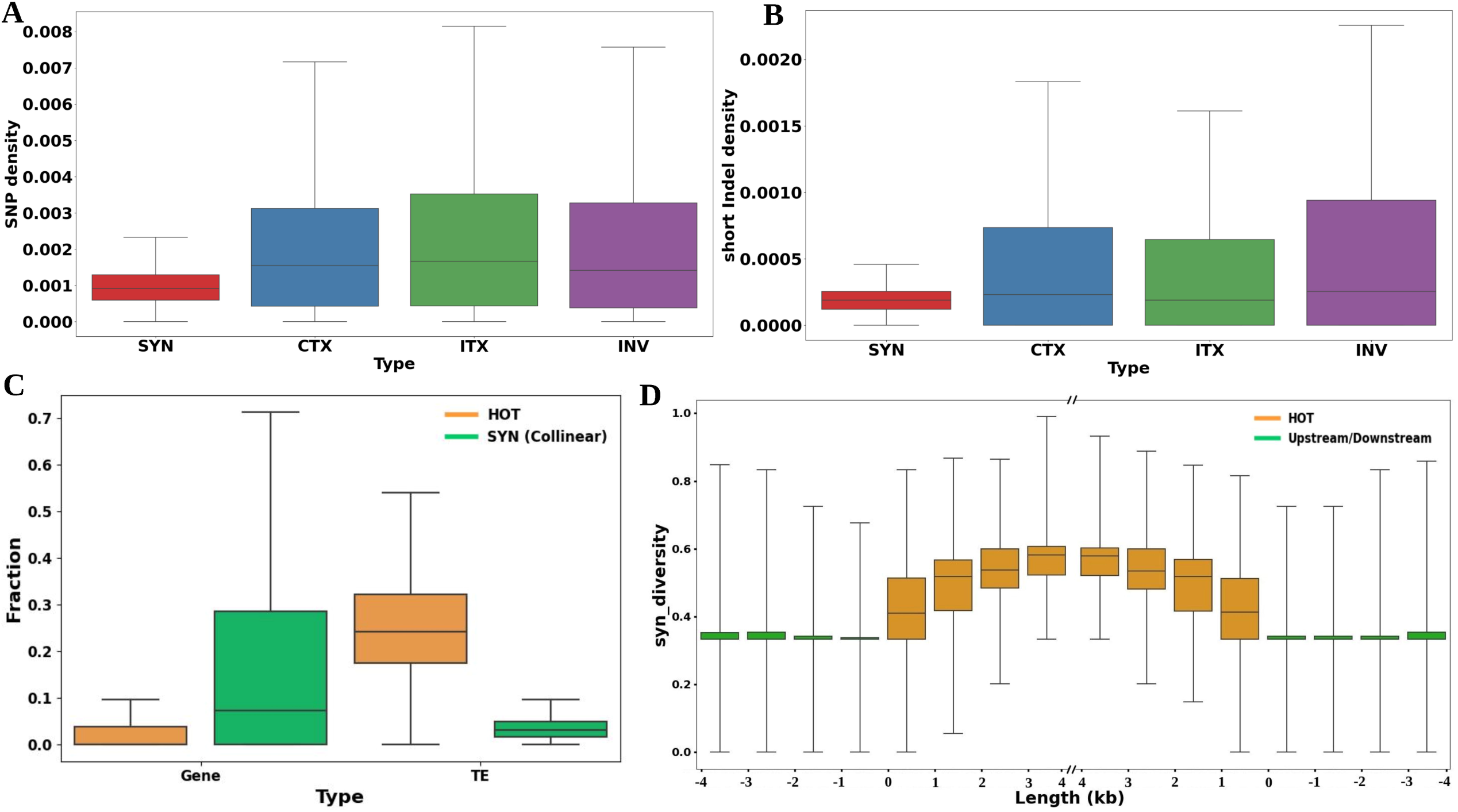
Sequence variation density and synteny diversity. **(A)** Density of single nucleotide polymorphisms (SNPs) per base pair (bp) in both syntenic and rearranged regions. **(B)** Frequency of short insertions and deletions (indels) per base pair in syntenic versus rearranged regions. **(C)** Gene and transposable element (TE) densities within 10,643 collinear regions (SYN) compared to 6,622 HOT regions. **(D)** The distribution of synteny diversity, measured in 1 kb sliding windows, around and within the 6,622 HOT regions.

### Quantification of genome collinearity and synteny diversity

The conservation of genome collinearity among different *desi* cattle has not been explored to date because of the absence of chromosome-level assemblies of different *desi* breeds. The chromosomal-level assemblies developed in this study provide an opportunity to investigate genome collinearity between breeds. We assessed collinearity using the synteny diversity parameter, π_syn_. Although π_syn_ can be calculated for any given region, it is important to establish collinearity within the context of whole genomes to avoid the misassignment of homologous but non-allelic regions.

Values of π_syn_ range from 0 to 1, with higher values indicating a complete absence of collinearity and lower values representing fully collinear regions. We calculated π_syn_ in 5-kb sliding windows across the genome using pairwise comparisons of all six accessions (**Additional file 2**). We identified 10,643 regions spanning 87.3 Mb across all breeds with a π_syn_ value of 0, indicating perfect conservation at the base pair level. However, these regions constituted a relatively small portion, approximately 3.2%, of the total genome (2,700 Mb). We identified 6,381 genes within these collinear regions. Given their complete collinearity across all breeds, these genes are likely to be conserved at the species level. To verify this assertion, we compared these genes to a set of 9,226 conserved orthologous genes from the BUSCO “mammalia_odb10” database. This analysis revealed a strong correlation between sequence-level collinearity and gene conservation, as ∼41% of the collinear genes (2,619) had corresponding matches with conserved mammalian orthologous genes. Conversely, rearranged regions exhibited higher π_syn_ values, with larger rearrangements, especially inversions and translocations, demonstrating elevated π_syn_. For instance, the largest inversion on chromosome X displayed an elevated π_syn_ value **(Figure 4B).**

### Hotspots of rearrangements

We further examined synteny diversity (π_syn_) across all chromosomes to identify regions prone to genomic rearrangements. Regions with higher π_syn_ values are enriched for complex rearrangements and diverse haplotypes. We defined hotspots of rearrangements (HOT regions) as genomic segments with a π_syn_ value exceeding 0.5. A total of 6,622 HOT regions, spanning 55.2 Mb, were identified across all chromosomes. Chromosome 10 exhibited the highest number of HOT regions (511), spanning a total of 4.7 Mb, which represents 4.5% of the chromosome. In contrast, chromosome 17 contained 129 HOT regions, spanning a total of 907 kb (**Supplementary Table S9**).

HOT regions were characterized by a higher proportion of TEs and a lower gene density compared to syntenic regions (**Figure 5C)**. Despite this, these regions contained a substantial number of genes (3,133), including protein-coding genes, long non-coding RNAs (lncRNAs), and other non-coding RNAs (**Supplementary Table S10**). To explore the potential functional implications of these genes, we performed GO and QTL analyses on the 2,794 identified protein-coding genes.

To investigate the conservation of synteny diversity in the vicinity of HOT regions, we examined the distribution of π_syn_ values in 1 kb sliding windows upstream and downstream of these regions. Despite the presence of rearrangements within HOT regions, the borders of these regions were well-conserved and exhibited similar patterns of synteny diversity on either side (**Figure 5D**).

### Functional characterization of genes in HOT regions

To elucidate the functional implications of protein-coding genes within the identified HOT regions, we performed GO annotation analysis. Of the 2,794 protein-coding genes, a substantial proportion was classified into biological processes (BP), molecular functions (MF), and cellular components (CC). Within the BP category, genes involved in immunological responses were significantly enriched, including antigen processing and presentation of endogenous antigen (GO:0019883), antigen processing and presentation of endogenous peptide antigen via MHC class I (GO:0019885), regulation of T cell mediated cytotoxicity (GO:0001914). Additionally, genes were enriched with GO terms associated with cell adhesion (**Figure 6A)**. The CC analysis revealed enrichment in sarcolemma (GO:0042383) and actin cytoskeleton (GO:0015629) (**Supplementary Figure S2A**). Analysis of MF indicated enrichment for GTPase regulator activity (GO:0030695) and peptide antigen binding (GO:0042605) (**Supplementary Figure S2B**). The presence of a substantial number of immune-related genes in HOT regions highlights the potential regulatory role of these regions in immune responses and cellular functions.

**Figure 6:**
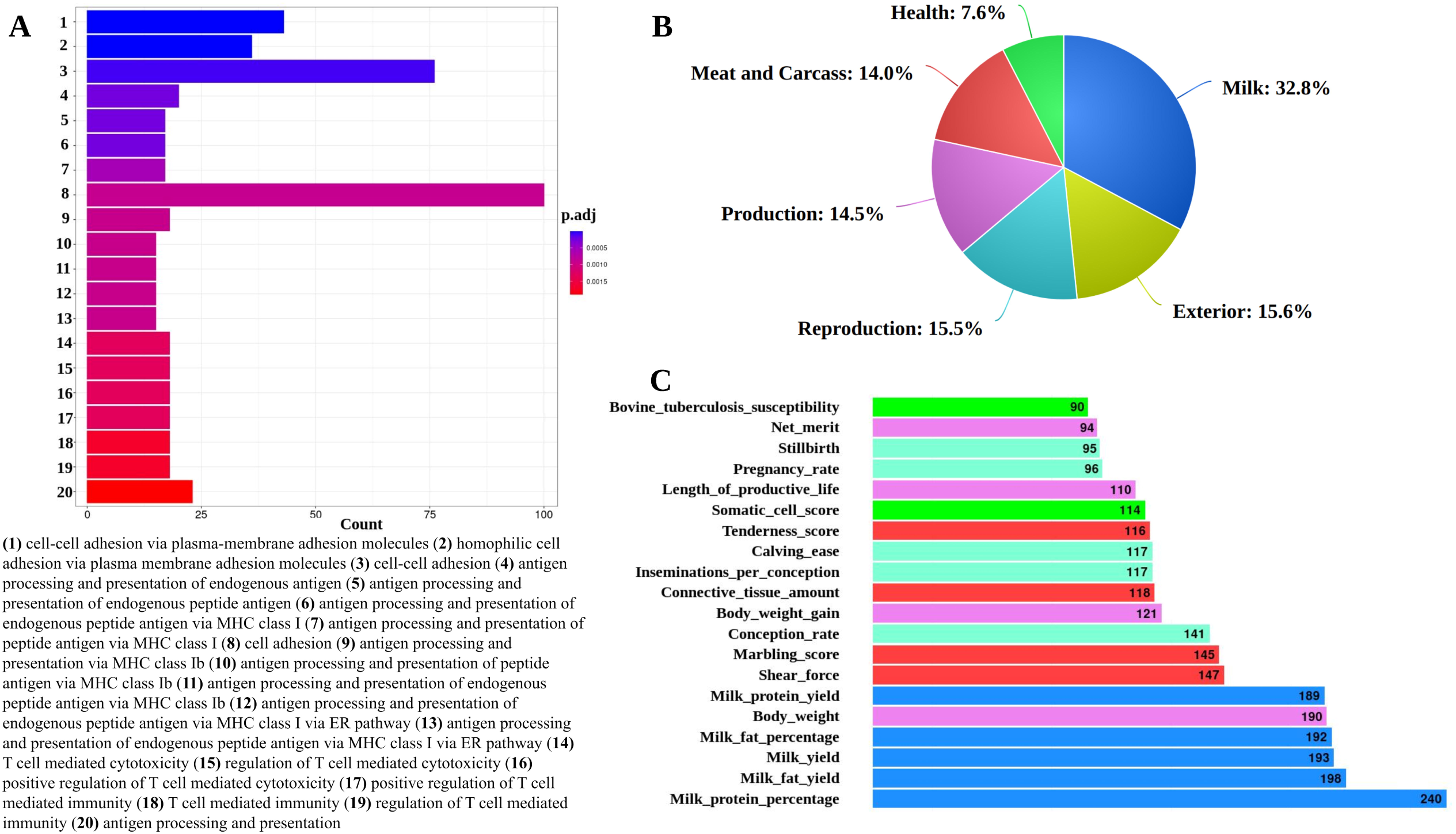
Functional annotation of genes in the HOT regions. **(A)** Bar plot illustrating the top 20 enriched GO terms for biological processes associated with genes in the HOT regions. **(B)** Pie chart representing the percentage distribution of different QTL trait types linked to genes within the HOT regions. **(C)** Bar plot displaying the top 20 enriched QTLs associated with various traits, with bar colors corresponding to the different trait categories as indicated in the pie chart (B) and count represents the number of genes associated with each QTL.

We further compared genes within HOT regions to putative QTL associated with a variety of traits. Our analysis revealed a notable enrichment of genes involved in milk production traits in HOT regions that overlapped with QTL regions. Of the 2,794 protein-coding genes in HOT regions, 1,644 (58%) were located within known QTL regions. Of these 1,644 overlapping genes Approximately 539 or 32.8% of genes within these overlapping regions are involved in milk-related traits (**Figure 6B)**. Furthermore, the most enriched QTLs encompassed not only milk production traits but also meat, carcass, and health traits, including bovine tuberculosis susceptibility and somatic cell score (**Figure 6C)**. These findings highlight the broader impact of HOT regions on the genetic architecture of complex traits relevant to both production and health.

### Synteny diversity in immune gene clusters of *desi* Cattle

We identified the loci of three major IGCs in the Brahman genome: Major Histocompatibility Complex (MHC) on chromosome 23 (29.40-29.65 Mb) (**Figure 7A)**, the Leukocyte Receptor Complex (LRC) on chromosome 18 (2.42-2.78 Mb) (**Supplementary Figure S3A**) and the Natural Killer Complex (NKC) on chromosome 5 (20.81-21.07 Mb) (**Supplementary Figure S4A**). MHC molecules are critical for antigen presentation, allowing the immune system to distinguish between self and non-self. In cattle, MHC genes are highly polymorphic and divided into Class I and Class II, where Class I molecules present intracellular antigens and Class II present extracellular antigens. The LRC, located on chromosome 18, encodes immunoglobulin-like receptors, which modulate immune cell activation and inhibition, particularly affecting NK cells and T cells. Similarly, the NKC, located on chromosome 5, encodes C-type lectin-like receptors that regulate NK cell-mediated recognition of MHC Class I molecules, crucial for immune surveillance against infected or abnormal cells. Gene copy numbers within these immune clusters varied across breeds (**Figure 7B**; **Supplementary Figure S3B**; **Supplementary Figure S4B**), which can impact immune function by influencing gene expression levels and receptor diversity. A scan of synteny diversity across these IGCs revealed that they coincide with the previously identified HOT regions on their respective chromosomes (**Figure 7C**; **Supplementary Figure S3C**; **Supplementary Figure S4C**), suggesting the functional importance of the HOT region in immune diversity within cattle breeds.

**Figure 7:**
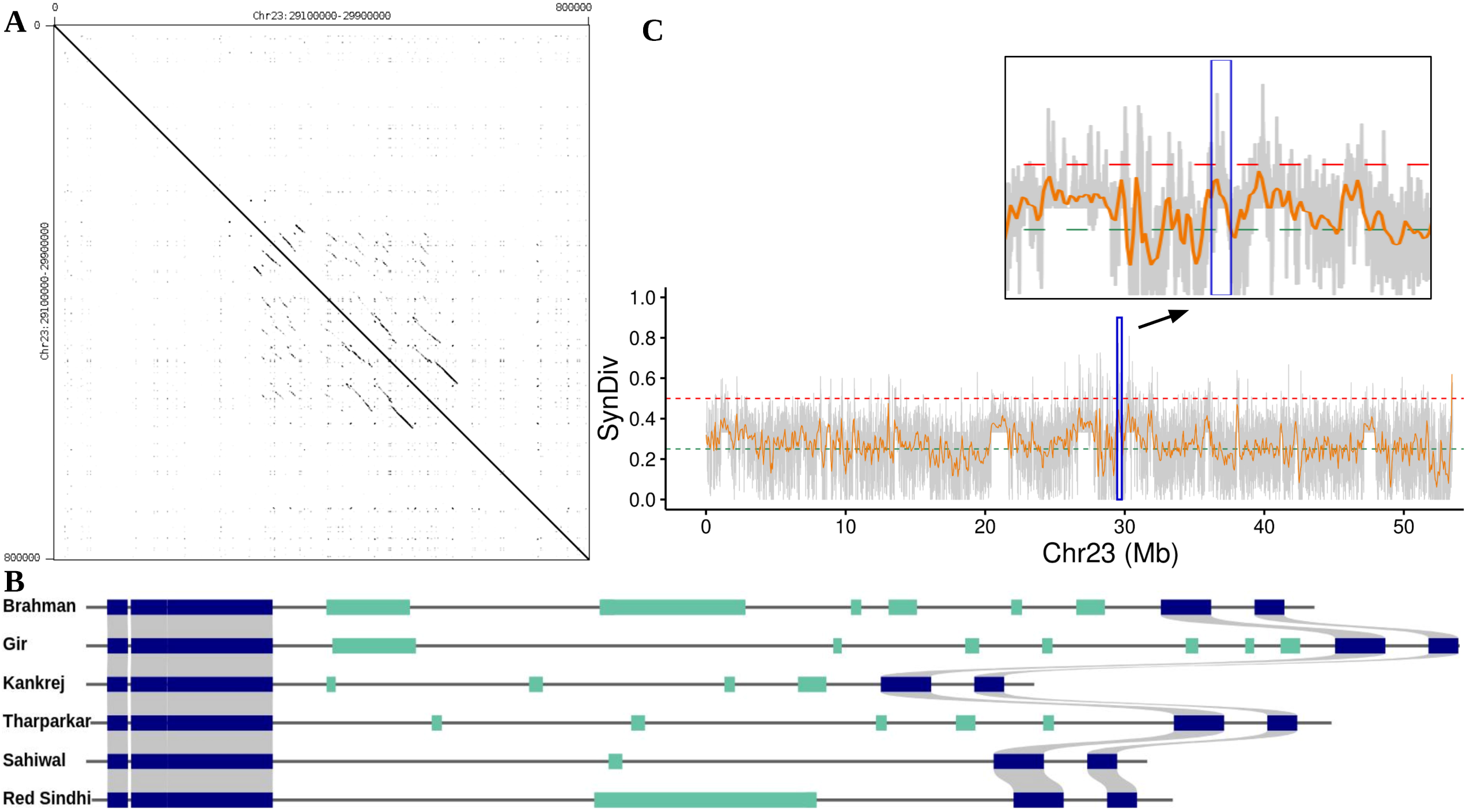
MHC cluster in the HOT regions. **(A)** Dot plot illustrating the MHC cluster region within the Brahman genome. **(B)** The figure depicts annotated protein-coding genes in MHC cluster as colored rectangles: blue rectangles represent non-immune related genes, and green rectangles represent immune-related genes. Grey lines connecting the rectangles indicate homologous relationships between non-immune genes. **(C)** Synteny diversity along chromosome 23, analyzed using sliding windows: orange line represents 100 kb sliding windows with a 50 kb step size, while the grey background shows 5 kb sliding windows with a 1 kb step size. Dashed green and red lines mark the thresholds for synteny diversity values at 0.25 and 0.50, respectively. The zoomed-in section highlights the synteny diversity specifically within the MHC cluster region on chromosome 23.

## Discussion

The *de novo* genome assembly of five *desi* cattle breeds: Gir, Kankrej, Red Sindhi, Sahiwal and Tharparkar using linked-reads 10X Chromium technology represents a significant advancement in the genetic characterization of these valuable livestock resources. This study successfully constructed highly contiguous assemblies for each breed, demonstrating the effectiveness of the linked read sequencing approach in assembling large, complex genomes, as has been previously shown in studies of humans and other species (34–37). While all five breeds resulted in high-quality assemblies, Sahiwal and Red Sindhi consistently outperformed the other breeds in measures of contiguity and scaffold length at each step of *de novo* assembly. This disparity can be attributed to such factors as library preparation quality and optimization of molecule length (53), and this suboptimal processing can lead to a more fragmented Supernova assembly, as observed in Gir, Kankrej and Tharparkar. These findings underscore the importance of meticulous optimization of sequencing protocols to achieve optimal assembly results (54, 55). To further enhance the quality of the assemblies, we employed the ARCS tool to improve the Supernova assemblies for each breed. This approach has been previously demonstrated to be effective in improving Supernova assemblies in other genomes (56, 57). In our case, ARCS was particularly effective in refining the more fragmented assemblies of Gir, Kankrej and Tharparkar. The resulting assemblies, especially for Sahiwal and Red Sindhi, exhibited high contiguity and large N50 values, indicative of high-quality genomes (58, 59). The successful assembly of 90% of the Sahiwal and Red Sindhi genomes into just 56 and 86 scaffolds, respectively, demonstrates the usability of linked-read technology in generating contiguous assemblies approaching chromosomal length. These results are comparable to the *Bos indicus* reference genome (54, 60), highlighting the technology’s potential for producing high-quality reference genomes for livestock species. Reference-guided assembly further improved the completeness of these genomes, with up to 96.7% of the genome assembled into pseudomolecules (58, 61). This result underscores the value of leveraging closely related reference genomes to enhance assembly quality and facilitate downstream analyses.

The completeness of the genome assemblies was assessed using BUSCO, which evaluates the presence of universal single-copy orthologs (BUSCOs) (59, 62). The high BUSCO scores, ranging from 93.3% to 95.4% for each genome, comparable to the Brahman reference genome (95.7%) indicate that the assemblies are nearly complete and highly accurate. These high scores also suggest that the genomes are highly representative of the functional gene content expected for these breeds. The annotation process revealed a large core set of shared genes across the breeds, along with unique genes. Observed variations in gene copy number, gene loss, or gene gain across the breeds are likely a result of genomic rearrangements, such as deletions, duplications, and inversions mechanisms known to drive genetic diversity and evolution within and between species (63, 64). Additionally, the identification of various non-coding RNAs enhances the functional annotation, offering a deeper understanding of the regulatory mechanisms that govern key phenotypes (60, 61). The consistent distribution of repeat elements across the assemblies, constituting approximately 47% of the genome, aligns with known patterns in other mammals. In Brahman cattle, around 49% of the genome consists of repeat elements, which is comparable to other mammalian assemblies, including human (GRCh38), Hereford cattle (ARS-UCD1.2), water buffalo (UOA_WB_1), and goat (ARS1) (26). These TEs play a crucial role in genome evolution (54, 58). The prevalence of LINEs, SINEs, LTRs, and DNA elements mirror the typical mammalian repeat landscapes that are essential for understanding genome plasticity and evolutionary processes (60, 61).

This study delves into the complex genomic architecture of five important Indian cattle breeds, revealing crucial insights into levels and characteristics of synteny, rearrangements, and breed-specificity. The similarities and differences in these high-quality assemblies will begin to ascribe their genomic implications. Detecting synteny in these breeds is particularly challenging because of gene loss, duplications, transpositions, and chromosomal rearrangements, all of which can introduce artifacts and complicate comparisons among genomes (65, 66). Each genome shows a high degree of synteny (87%-95%) against the Brahman reference genome, suggesting a significant shared genomic architecture.

This result is expected, as similar levels of synteny have been observed in humans when comparing two complete telomere-to-telomere (T2T) genomes, T2T-CHM13 from northern European descent (67) and T2T-YAO from Chinese origin (20). These assemblies share a 95.32% syntenic region (**Supplementary Table S11**). Likewise, wild goats (*Capra ibex*) show 90% synteny with the domestic goat (*Capra hircus*) genome (68). The high level of synteny observed among these breeds of cattle underscores their shared genetic similarity (69, 70). The contiguity of genome assembly plays a crucial role in the accurate detection of SVs, further highlighting the importance of high-quality assemblies in genomic studies. Therefore, a higher degree of synteny was detected between Red Sindhi and Sahiwal, among breeds, likely because the genome of these breeds had the highest levels of contiguity among the five genome assemblies.

Large structural variations, such as inversions, translocations, and duplications, are known to influence gene expression, genomic stability, and phenotypic diversity (61). Our catalog of SVs in desi cattle comprises thousands of variants, highlighting their potential to significantly influence phenotypic traits. Further studies will need to be done to determine the impacts of these SVs. One example worth investigating is the inversion on chromosome 12, found across all the breeds against Brahman, which includes the LMO7 gene. This gene is crucial for both skeletal and cardiac muscle development and is highly conserved across vertebrates, from fish to mammals (71, 72). Additionally, LMO7 has been implicated in macrophage function, where it plays a role in degrading PFKFB3 through K48-mediated ubiquitination, a process that modulates inflammatory responses, including the pathogenesis of inflammatory bowel disease (73). The presence of this inversion may contribute to both structural and functional variations within these cattle breeds, potentially influencing traits related to muscle development and immune function but, the impact of this inversion was outside the scope of this study.

We also identified 10,643 regions spanning 87.3 Mb with π_syn_=0, indicating perfect collinearity across all breeds. These regions, representing approximately 3.2% of the total genome, include 6,381 genes that are highly conserved across breeds. This finding highlights an important enrichment of genes within a relatively small portion of the genome, accounting for approximately one-third of the functional space. Our sequence-based synteny identification approach aligns with previous gene-based synteny studies in other species, confirming that collinear regions are enriched with genes (65, 70). Intersecting these 6,381 genes with the BUSCO “mammalia_odb10” database revealed a strong association between sequence-level collinearity and gene conservation, with ∼41% of these genes matching conserved mammalian orthologs. This finding underscores the evolutionary importance of conserved synteny in maintaining essential biological functions across species.

A key finding in our study is the identification of several regions with high synteny diversity (π_syn_ values exceeding 0.5) across the genomes. Using a 5 kb sliding window, we were able to map areas of genomic conservation and divergence. The regions of divergence, termed HOT regions, contain non-collinear haplotypes that are more likely to undergo or preserve complex mutations and do not spread through the population by traditional means such as meiotic recombination or haplotype-specific drift (17, 74). Instead, these regions may evolve rapidly due to the local accumulation of mutations that are not effectively reshuffled by recombination. We identified 6,622 HOT regions spanning a total of 55.2 Mb across all chromosomes. While the precise impact of these HOT regions remains unclear, their potential role in shaping genome architecture and function requires further investigation. Such rearrangements have been previously highlighted as crucial in driving evolutionary adaptation across various species (75).

Our analysis of HOT regions revealed a significant enrichment of immune-related genes, particularly within major IGCs such as the MHC, NKC, and LRC. The complete overlap of all IGCs within HOT regions is particularly intriguing, suggesting that these regions play a pivotal role in generating structural diversity essential for the adaptive immune response. The high variability in IGC gene copy numbers within HOT regions could significantly contribute to the production of a diverse repertoire of antibodies, which is vital for the immune function of these cattle. This observation is supported by earlier studies emphasizing the importance of IGCs in evolutionary adaptation, particularly in response to environmental pressures and pathogens (76). Moreover, our GO and QTL analyses revealed significant enrichment of genes involved in immune functions, including those associated with antigen processing and T cell-mediated cytotoxicity. Notably, the overlap of HOT regions with enriched QTLs associated with Bovine tuberculosis susceptibility is particularly significant. For instance, the IL21R gene, located within these regions, is linked to the Bovine tuberculosis susceptibility QTL and is also enriched in GO terms such as “immune effector process” (GO:0002252) and “adaptive immune response” (GO:0002250). This gene has been reported to play a major role in the immune response against *Mycobacterium* spp. (77). These findings further underscore the potential regulatory role of HOT regions in immune responses, highlighting their importance in the cattle’s ability to adapt to disease challenges. Further studies are needed to elucidate the precise mechanisms underlying the association between HOT regions and IGC diversity, as well as to explore their potential role in other complex traits.

The overlap between HOT regions and QTL associated with economically important traits, such as milk production and health-related traits, further indicate the potential functional significance of these genomic regions. The identification of genes within HOT regions associated with milk production suggests that these regions may be critical for regulating traits vital for the economic viability and productivity of these cattle breeds.

## Conclusion

In this study, we successfully generated and analyzed high-quality *de novo* genome assemblies for five major *desi* cattle breeds, achieving substantial completeness and contiguity. For the first time in *desi* cattle, synteny and structural rearrangements at the genomic sequence level were comprehensively examined, revealing extensive syntenic regions with the Brahman reference genome, alongside significant structural variations, including large insertions, inversions, translocations, and duplications. Our analysis also demonstrated a higher prevalence of SNPs and Indels in rearranged regions compared to syntenic regions. Additionally, this study represents the first application of the synteny diversity (π_syn_) parameter for the comparative analysis of *desi* cattle, identifying both highly collinear regions and hotspots of genomic rearrangements. Conserved genes, such as universal genes, were predominantly located in collinear regions, while HOT regions were enriched for immune-related genes and functions. The discovery of IGCs within these HOT regions underscores their potential role in disease resistance and adaptation. These findings provide valuable insights into the genomic architecture of *desi* cattle breeds, marking a significant advancement in genomic research for these populations. This work opens new avenues for future research, breeding programs, and the improvement of cattle productivity and health.

## Data availability

The Linked-Read raw data was submitted to the Indian Biological Data Centre (IBDC) with the INDA study id INRP000127 (**Supplementary Table S12**). The specific accession numbers for the raw assembled genome can be found in **Supplementary Table S13**. All the data can also be accessed from NCBI/EMBL/DDBJ.

## Supplementary data

Supplementary tables are available at NAR online.

Supplementary figures are available at NAR online.

Additional file 1 are available at NAR online.

Additional file 2 are available at NAR online.

## Acknowledgments

The authors gratefully acknowledge the financial support provided by the Department of Biotechnology (DBT), Ministry of Science, New Delhi, India. We extend our sincere thanks to the National Institute of Animal Biotechnology (NIAB) for their invaluable support throughout the study. S.A. would also like to express gratitude to Dr. G. Taru Sharma, Director of NIAB, for her guidance and encouragement. M.N., C.P.V.T., and B.D.R. were supported by the appropriated project 8042-31000-112-000-D, “Accelerating Genetic Improvement of Ruminants Through Enhanced Genome Assembly, Annotation, and Selection” of the USDA Agricultural Research Service. Any mention of trade names or commercial products is solely for the purpose of providing specific information and does not imply recommendation or endorsement by the U.S. Department of Agriculture. The USDA is an equal opportunity provider and employer.

## Authors’ Contributions

S.A., R.K.G., S.N.R., and S.S.M. conceived and designed the study. S.S.M., R.K.G., and S.A. facilitated sample collection and sequencing. S.A. and B.D.R. assembled the genomes and performed the BUSCO analysis. S.A. and A.S. annotated the genomes for TEs, conducted Liftoff analysis, Syri analysis, IGC annotation, and collinear & hotspot region analysis. S.A. and A.S. drafted the manuscript. M.K. performed the Longranger analysis and GO analysis. M.N. conducted the QTL analysis. C.P.V.T., R.K.G., B.D.R., and S.N.R. edited the manuscript. All authors reviewed and approved the final manuscript before submission.

## Funding

Linked-Read raw data was generated as part of a research project funded by the Department of Biotechnology (DBT), India, entitled ‘Genomics for Conservation of Indigenous Cattle Breeds and for Enhancing Milk Yield, Phase-I’ (BT/PR26466/AAQ/1/704/2017).

## Competing Interest

The authors declare that they have no conflicts of interest.

## Notes

### Competing Interest Statement

The authors have declared no competing interest.

