## Supplementary Figures for "Genome assemblies of Indian *desi* cattle reveals hotspots of rearrangements and immune-related genetic diversity": Supplementary Figures.docx

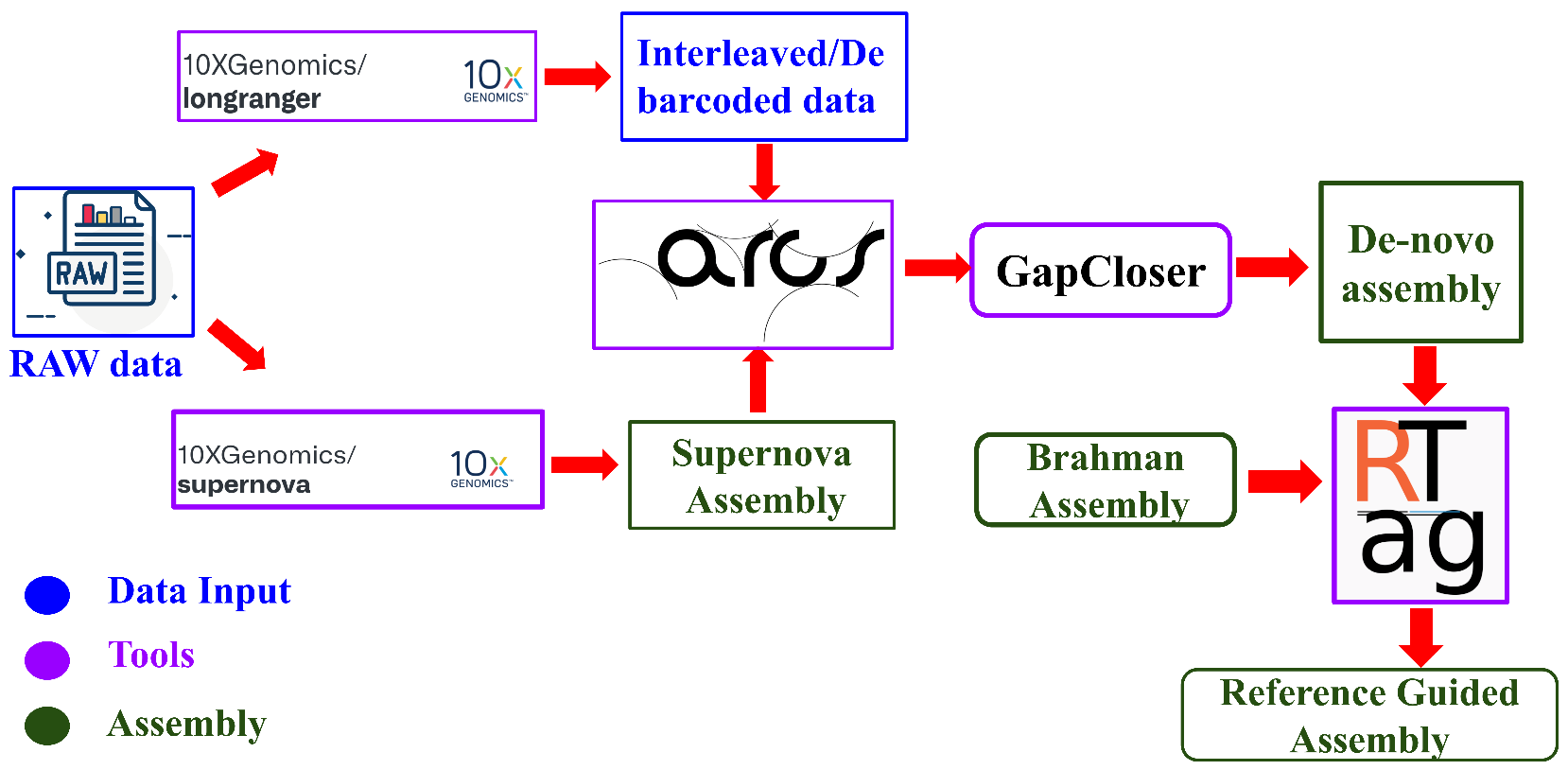

**Figure S1: Schematic workflow for the making of reference-guided genome assembly.**

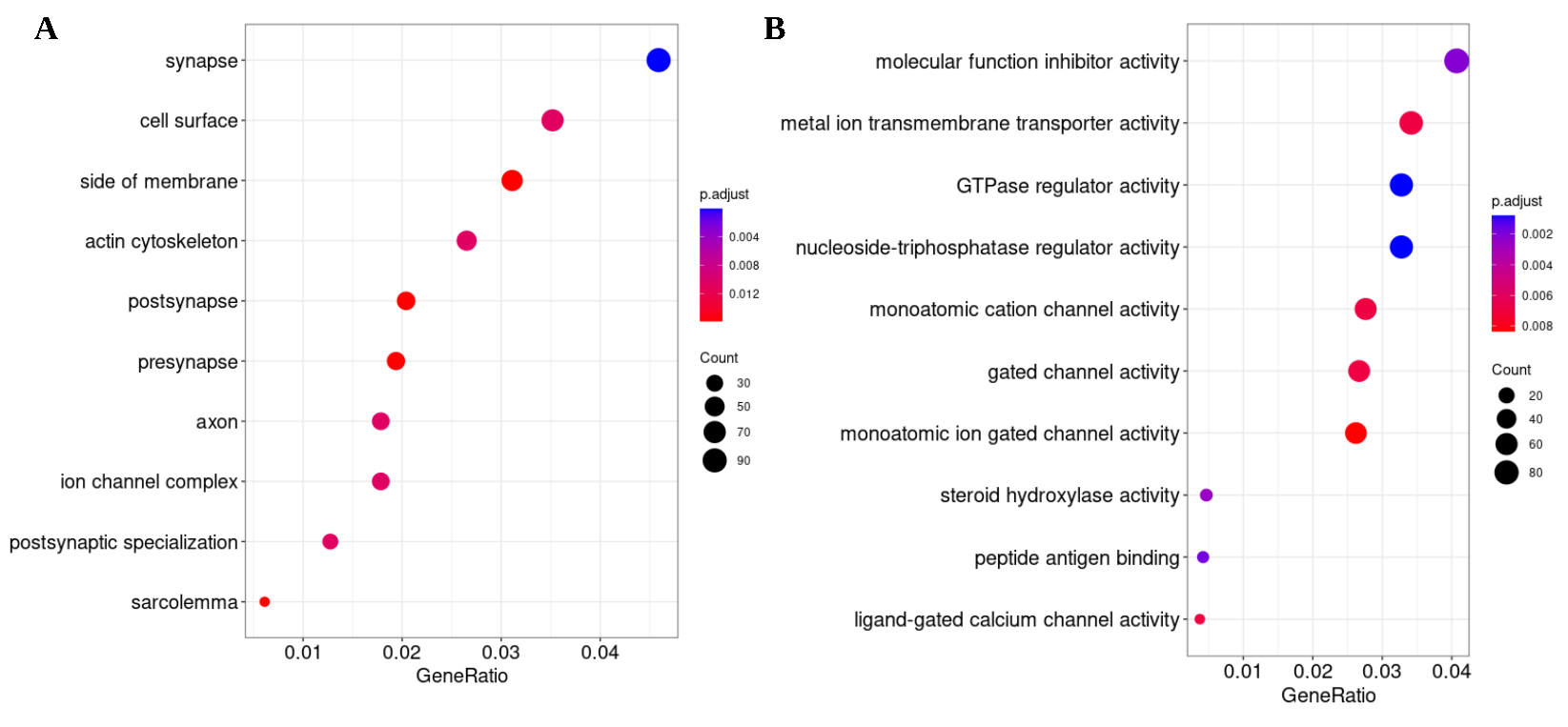

**Figure S2 : GO annotation of genes in HOT regions.(A)** Dot plot showing the top 10 enriched GO terms related to cellular components (CC) for genes located in HOT regions. **(B)** Dot plot depicting the top 10 enriched GO terms associated with molecular functions (MF) for these genes.

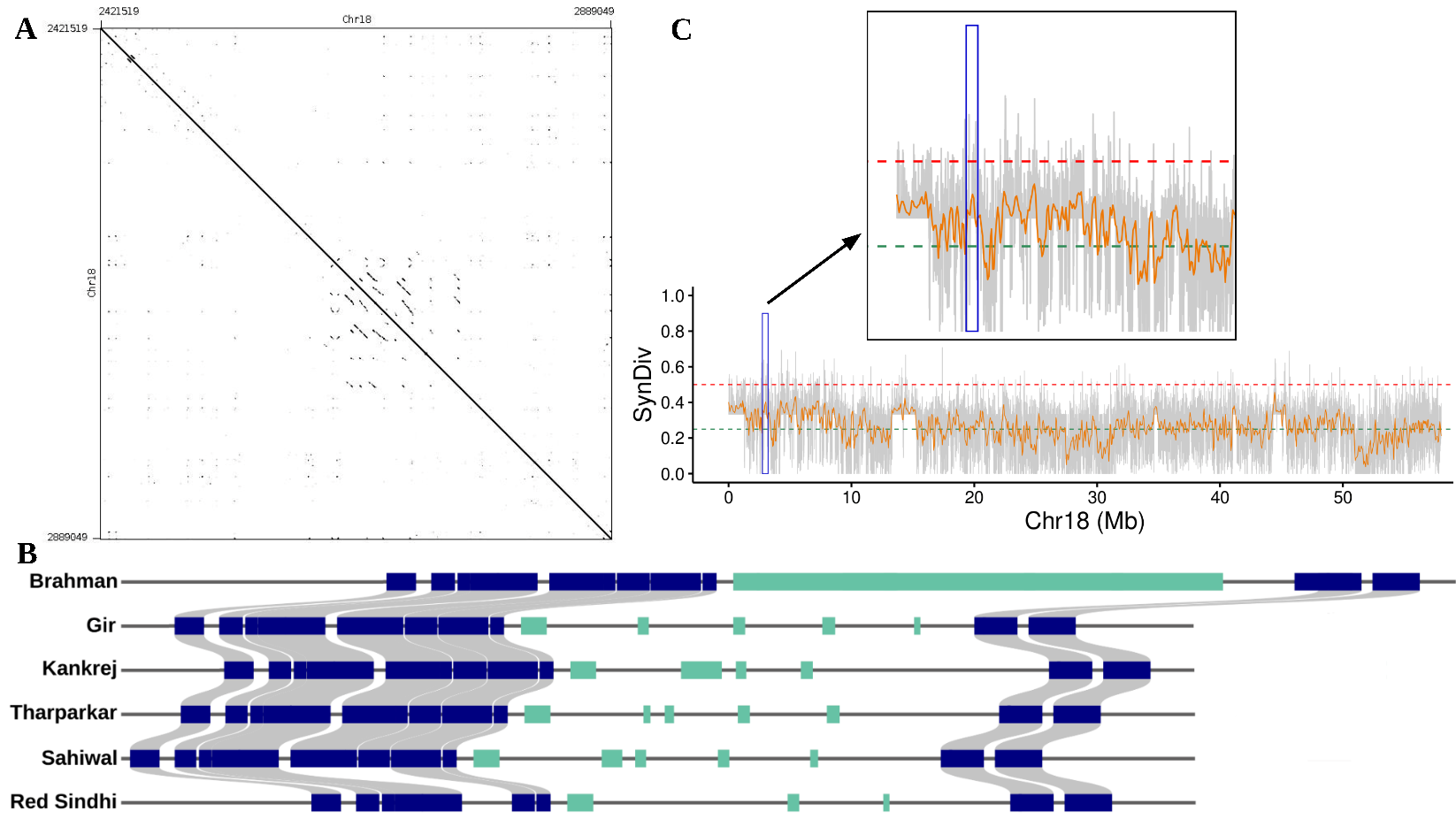

**Figure S3 : LRC cluster in the HOT region. (A)** Dotplot showing LRC cluster in chromosome 18. **(B)** The figure shows the annotated protein-coding genes in the LRC cluster, represented as colored rectangles: blue rectangles correspond to non-immune related genes, while green rectangles indicate immune-related genes. Grey lines connecting the rectangles indicate homologous relationships between non-immune genes. **(C)** Synteny diversity along chromosome 18, with zoomed-in view focusing on the LRC cluster region to highlight synteny diversity specifically within this area.

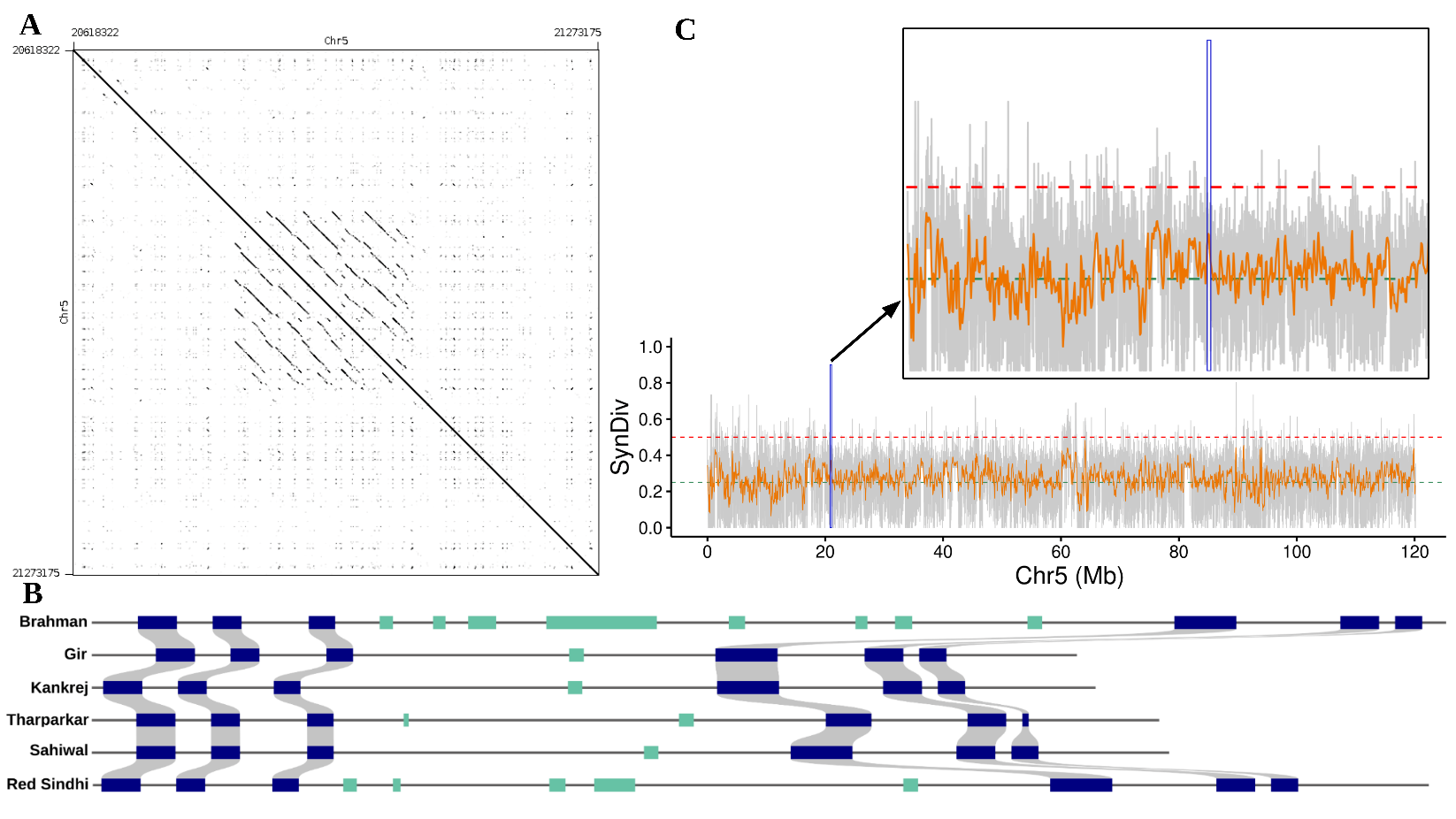

**Figure S4 : NKC cluster in the HOT region.** Legend same as (supplementary figure S3) in chromosome 5.
