## Supplementary figures and images for "Genome assemblies of Indian *desi* cattle reveals hotspots of rearrangements and immune-related genetic diversity"

### Additional_file2.pdf

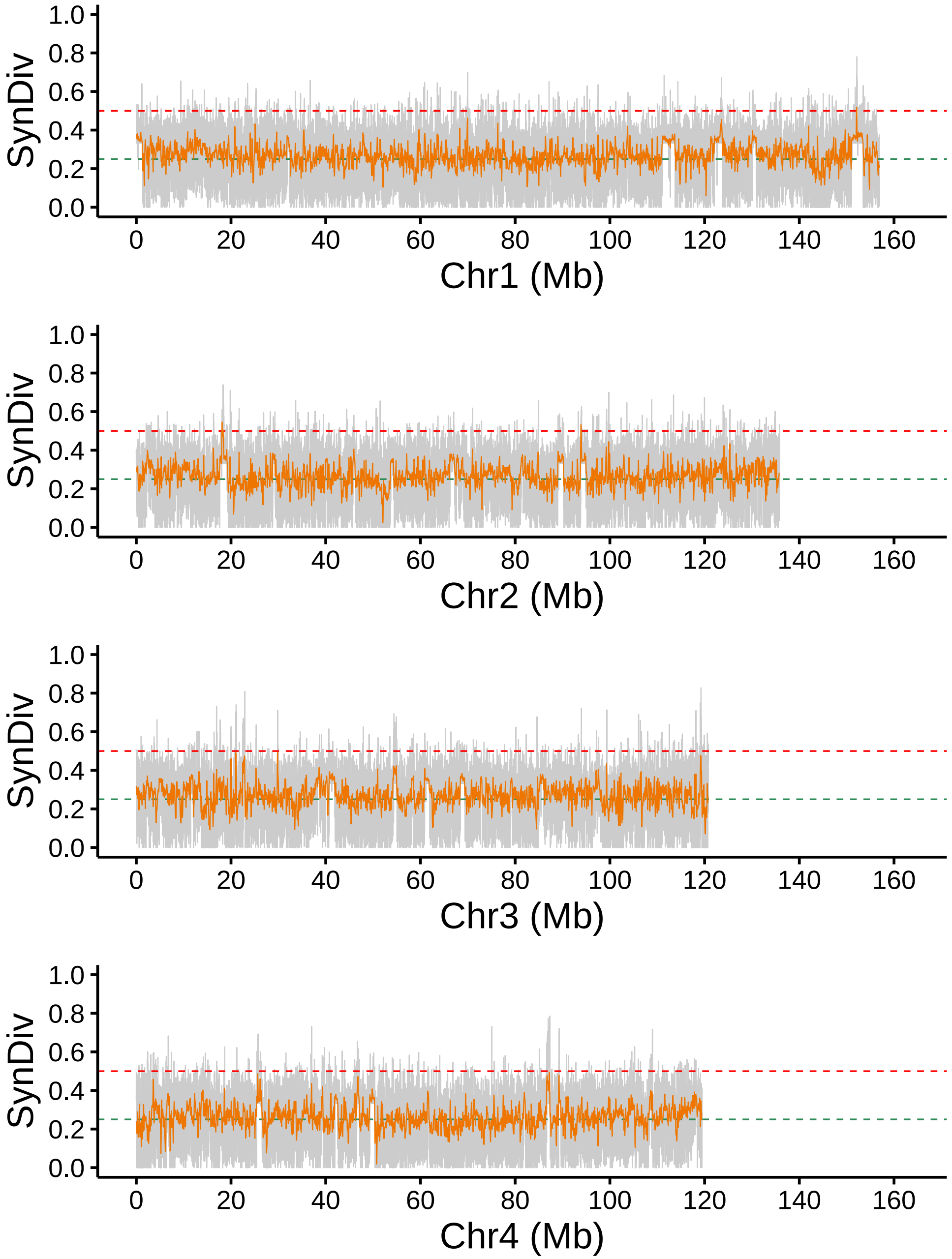

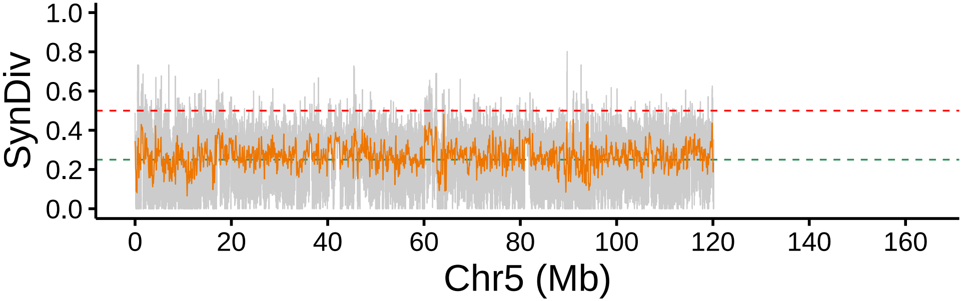

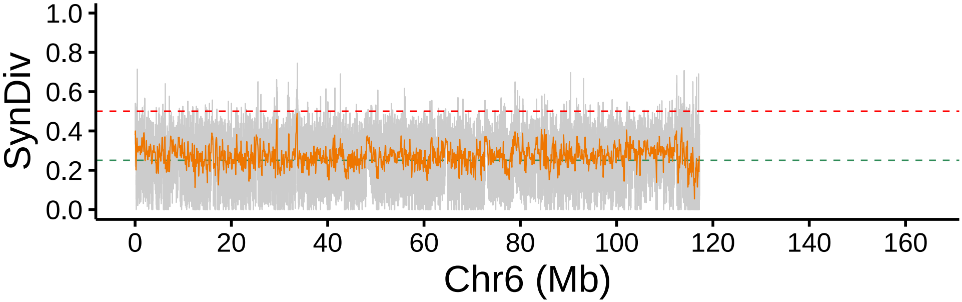

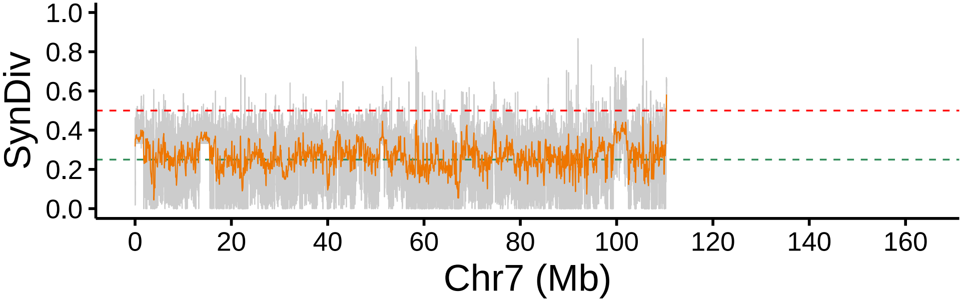

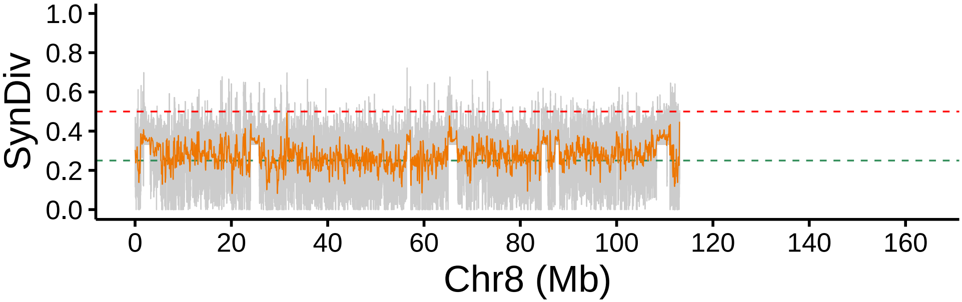

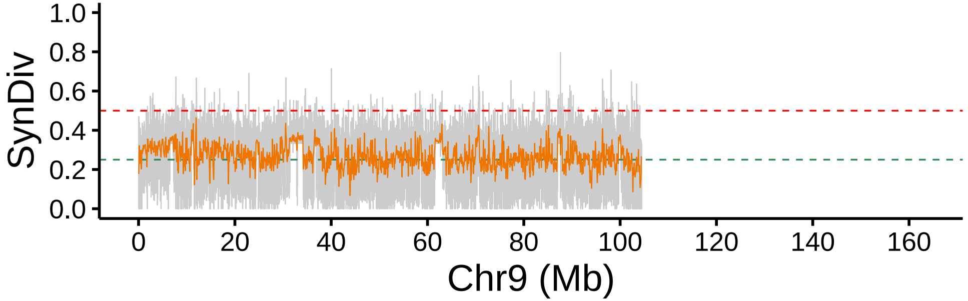

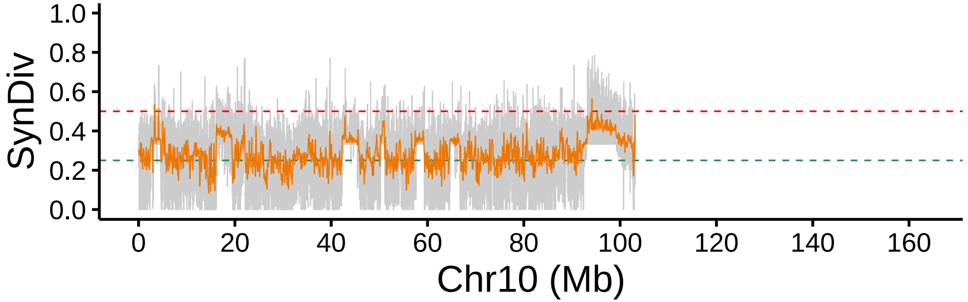

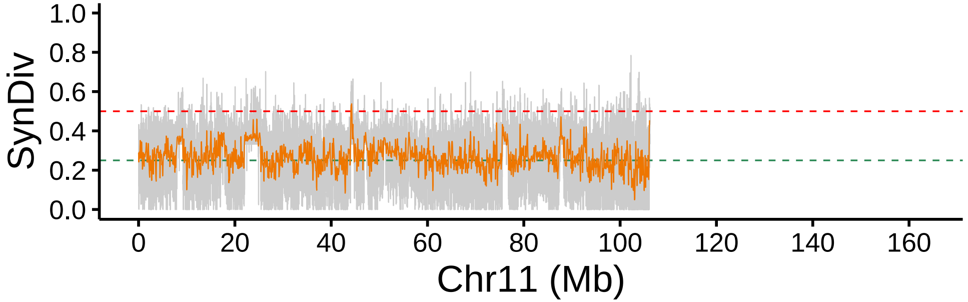

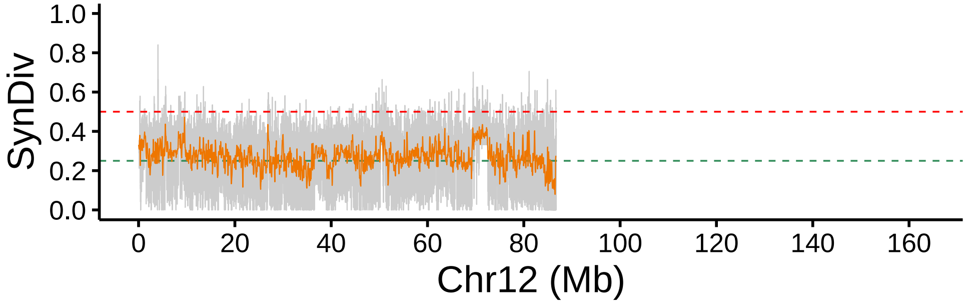

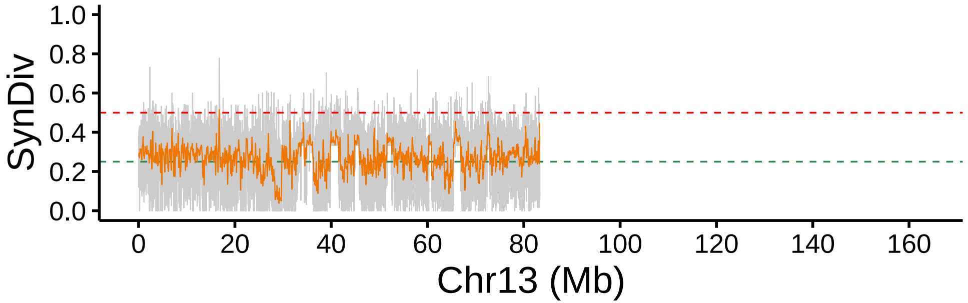

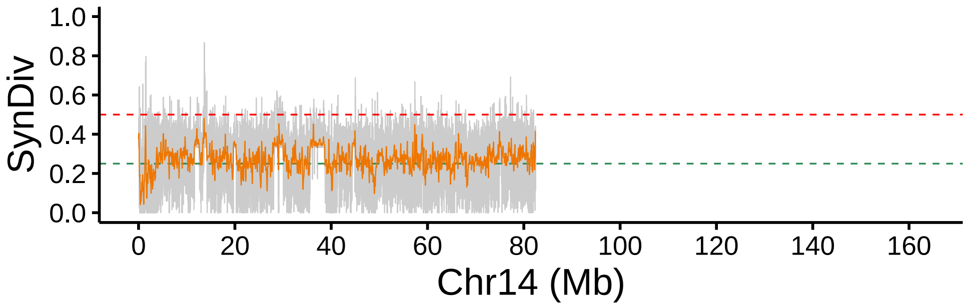

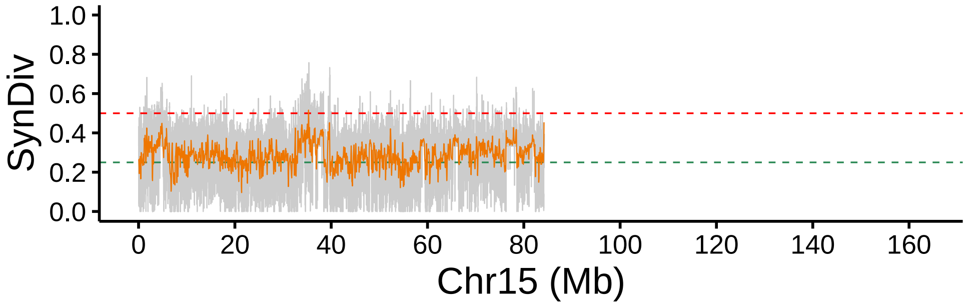

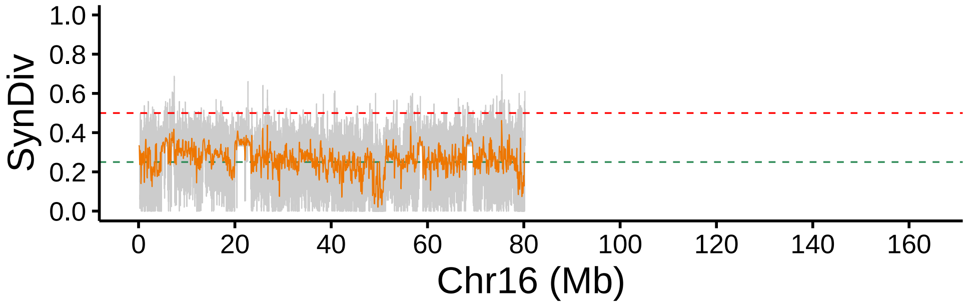

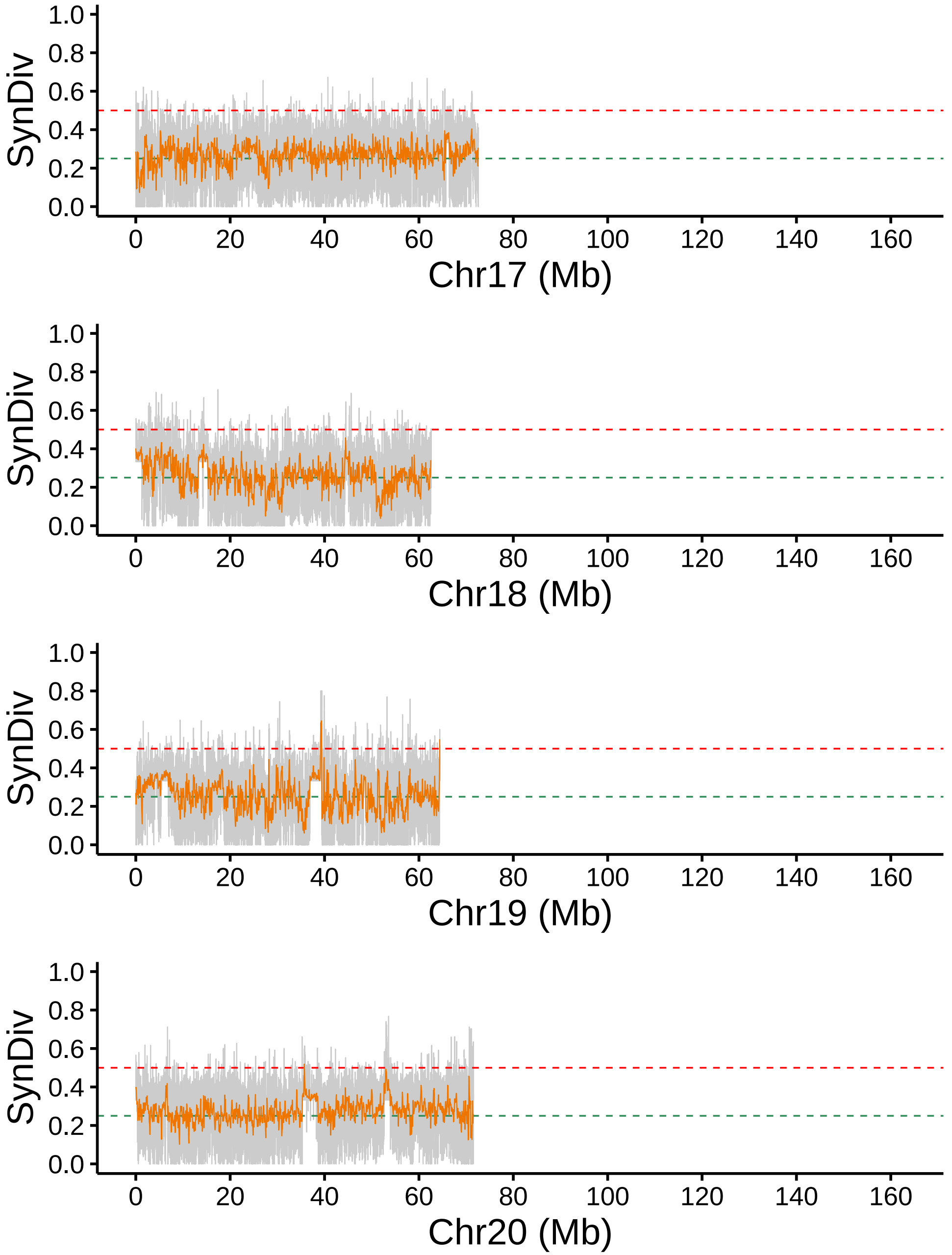

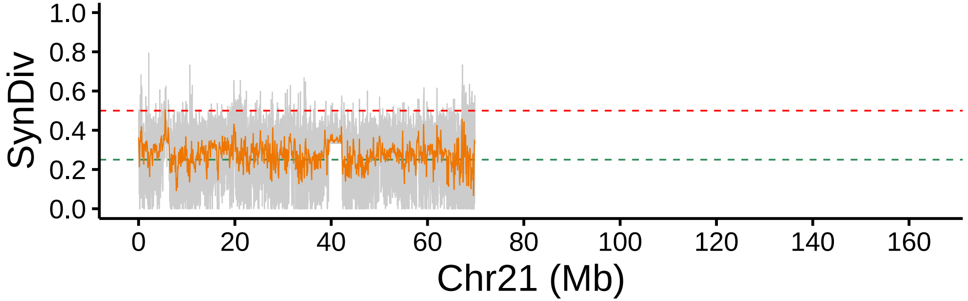

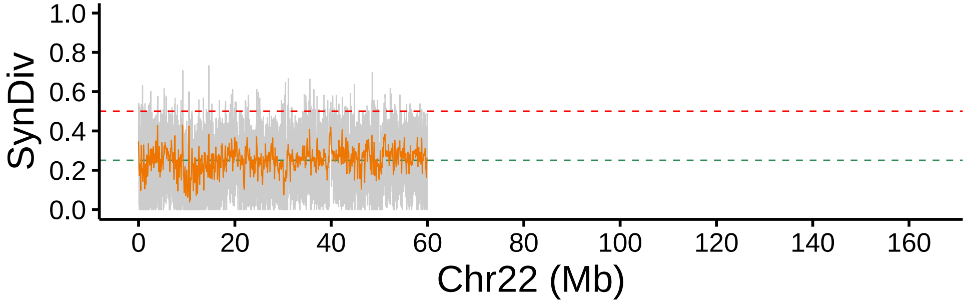

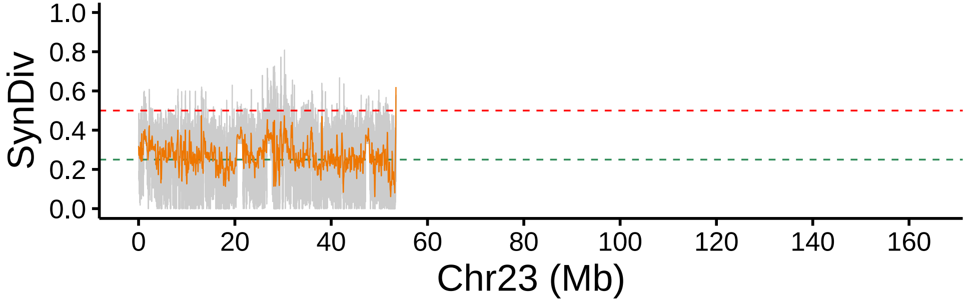

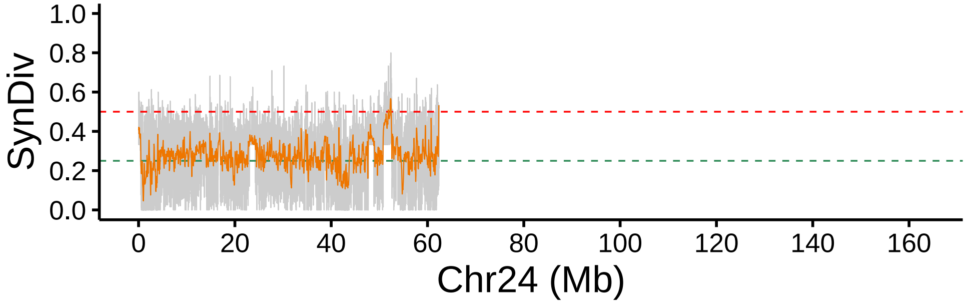

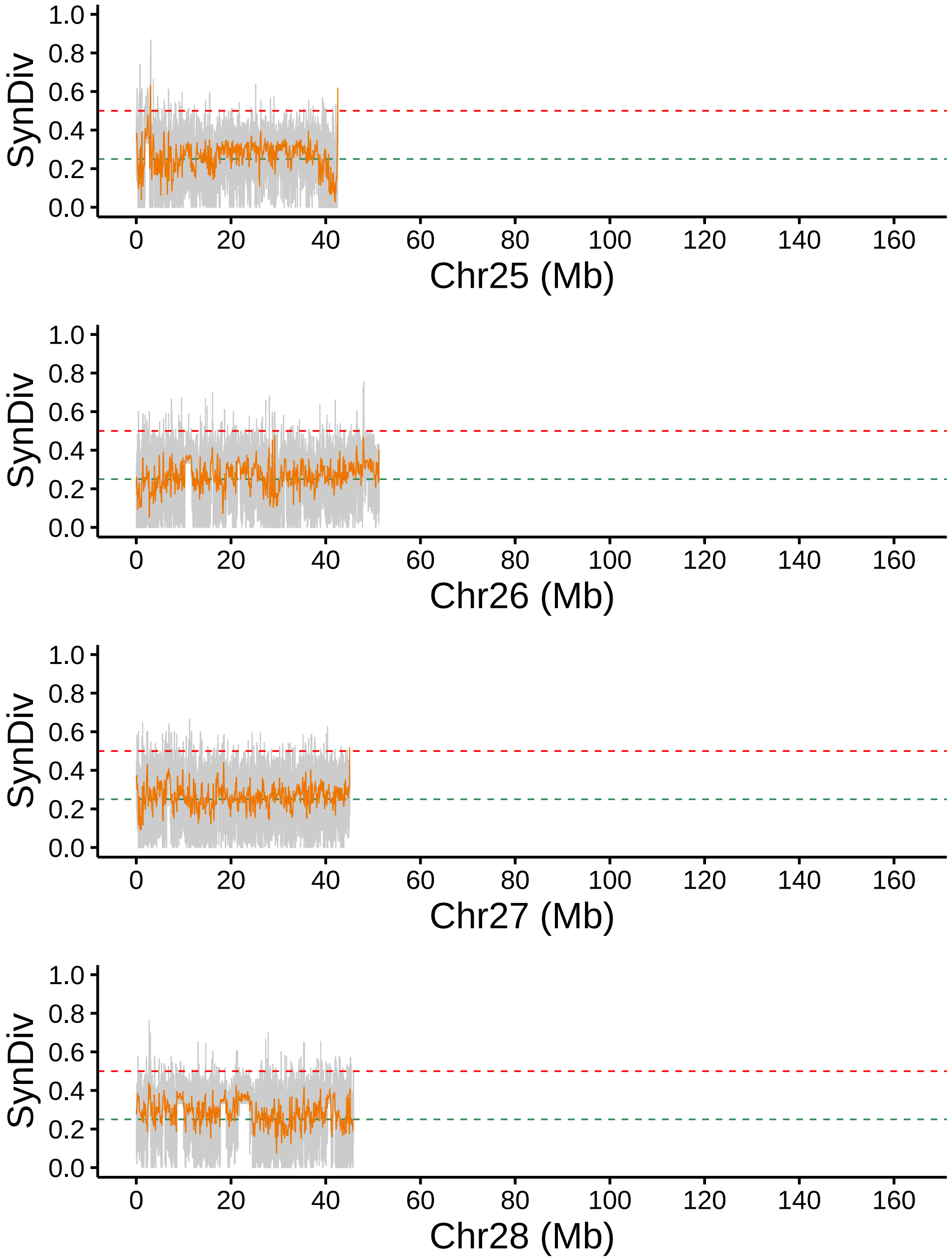

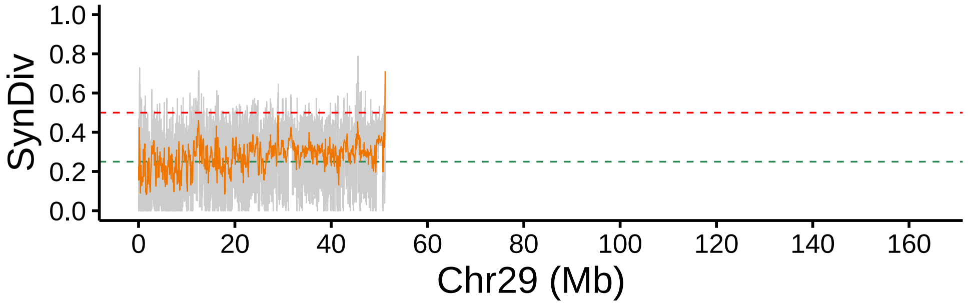

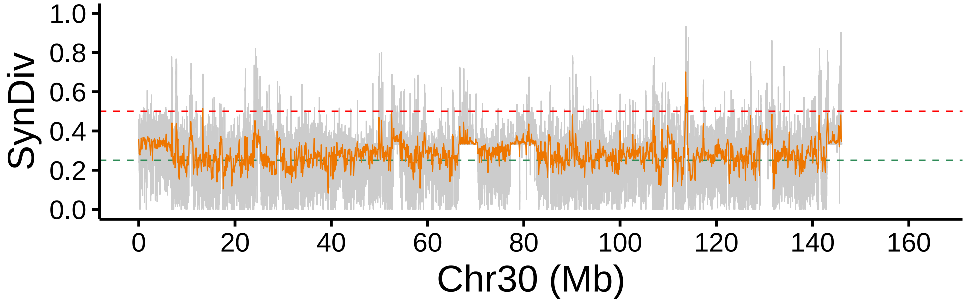
